# Structural modeling and experimental validation define the MxA-Thogotovirus nucleoprotein interface that drives restriction and escape

**DOI:** 10.64898/2026.08.07.743570

**Authors:** L. América Chi, Megan E. Levy, Rechel A. Geiger, Harmit S. Malik, Jagdish Suresh Patel

**Author notes:** Correspondence to be addressed to: Jagdish Suresh Patel, 875 Perimeter Dr MS 0904, Department of Chemical and Biological Engineering, University of Idaho, Moscow, ID, 83844, USA. These authors contributed equally and are listed alphabetically (L. América Chi, Megan E. Levy).

## Abstract

The human MxA (myxovirus resistance protein A) host restriction factor inhibits orthomyxoviruses, such as Thogotovirus (THOV) and influenza A virus (IAV), by binding to their nucleoproteins. Despite being discovered over six decades ago, how MxA interacts with viral targets remains unclear. Earlier studies using evolutionary analysis and mutagenesis showed that the MxA L4 loop, especially a hydrophobic aromatic amino acid at residue 561, is crucial for binding THOV nucleoprotein (NP) and restricting THOV. Here, we combined previous insights with structure prediction methods, molecular dynamics simulations, and experimental validation to define the human MxA L4 loop binding interface to THOV NP. We also evaluated the stability of MxA L4-NP binding through classical all-atom molecular dynamics simulations. Our model revealed MxA L4 binding to a surface-exposed site on THOV NP, including residues previously linked to viral escape from MxA restriction, even though this information was not used to guide our modeling efforts. This MxA-THOV NP interface is distinct from NP’s RNA-binding or oligomerization surfaces. Our molecular dynamics simulations also agree with earlier data indicating that F561Y enhances MxA binding to THOV NP, whereas F561W reduces it and F561V ablates it entirely. Based on this model, we predicted specific variants in human MxA or THOV NP that could result in increased host restriction or viral escape. We tested these predictions using a viral minireplicon assay to validate our model. Our efforts will guide vital viral surveillance studies and the development of MxA-based antivirals. *(240)*

**Significance Statement:** The interferon-stimulated MxA protein encodes a critical barrier to zoonotic spillover of orthomyxoviruses, restricting infection by binding viral nucleoprotein (NP). Despite being among the best-studied interferon-stimulated genes (ISGs), the specific host-viral interaction surface involved in binding or restriction is poorly understood. We address this hurdle using structural modeling and molecular dynamics simulations to generate a model of the biochemical interactions between MxA and Thogotovirus NP. Our model is consistent with previous work identifying mutations that strengthen or weaken the binding interaction. We validate this model using *in vitro* analysis of predicted mutations at the host-virus interface. Our model provides a framework for additional mutational studies, surveillance of viruses poised for zoonosis, and rational antiviral design. *(115)*

## Introduction

Orthomyxoviruses are negative-sense, segmented, single-stranded RNA viruses that pose a significant burden to public health, especially influenza viruses. Due to their high mutation rate and propensity for genetic reassortment^1,2^, novel influenza strains derived from those that usually circulate in other species, such as waterfowl and pigs, continue to acquire the ability to infect humans efficiently. This leads to a high risk of future pandemics; the recent rise in H5N1 cases in North American cattle populations and sporadic infections in humans are examples of the persistent risk imposed by such viruses^3^. Although influenza strains have been the primary focus of scientific attention, they are not the only orthomyxoviruses of concern. Thogotovirus (THOV) is a tick-borne orthomyxovirus that replicates in rodents^4,5^, but causes only sporadic cases of clinical infection in humans^6^. Viruses in the genus *Thogotoviridae* from Africa, Asia, Europe, and North America pose a significant risk of zoonotic spillover. For example, the Jos strain (JOSV) of Thogotovirus, identified in Nigeria, can infect humans^7^. Similarly, the recently described Bourbon virus from this genus has caused multiple deaths in the United States^8,9^.

One of the primary barriers to spillover of orthomyxoviruses into humans is MxA^10^. The Mx family of genes (myxovirus resistance) was first discovered in 1962 in pioneering studies on influenza-resistant mice^11^: the mouse *Mx1* gene conferred complete protection against lethal doses of influenza A virus in some mouse strains, whereas strains lacking Mx1 were highly susceptible^11,12^. *Mx1* was later determined to be an interferon-responsive gene^13^ that encodes a dynamin-like GTPase^14–16^. Humans encode two *Mx* genes^17^: *MX2* encodes the nuclear pore-associated MxB protein, which confers protection against retroviruses and herpesviruses^18–22^, whereas its paralog *MX1* encodes the cytoplasmic MxA protein, which has broad antiviral activity against a variety of viruses^23^, including *Orthomyxoviridae* (Influenza, Thogotovirus), *Bunyaviridae* (La Crosse virus^24–27^), and DNA viruses (Hepatitis B virus, African swine fever virus^28–31^). This broad activity is thought to be partially achieved through MxA’s dynamin-like GTP-dependent mechanism of oligomerization into ring structures that entrap viral nucleocapsids^32^.

Structural analyses have revealed three functional domains in MxA proteins: an N-terminal GTPase module, a central bundle signaling element, and a C-terminal stalk containing specificity-determining loops that recognize viral targets, such as nucleoproteins (NPs) from THOV or Influenza A virus (IAV)^33^. This mechanistic versatility allows MxA to disrupt multiple stages of viral replication, from nucleocapsid trafficking^34–36^ to genomic transcription^37^. Originally believed to be evolutionarily constrained to vertebrates, Mx proteins are now thought to have evolved in early eukaryotes^38^. Despite their prominent role in interferon-mediated protection from orthomyxovirus spillover^10^ and as broad-spectrum antiviral effectors^23^, the biochemical details of MxA interactions with any viral target remain unknown.

To restrict orthomyxoviruses, MxA targets NP oligomers^39^, which are a critical component of the viral ribonucleoprotein complex. Nucleoprotein complexes are involved in diverse functions of the orthomyxovirus life cycle, including transport, transcription, and packaging of the viral genome^40–42^. Although the specific binding modalities between MxA and NP remain unknown, MxA’s binding affinity for viral NP correlates with MxA’s antiviral restriction^43^, whereas the lack of binding is associated with viral escape from MxA^7^. Evolutionary analyses of natural and laboratory-generated MxA restrictor variants, as well as orthomyxoviruses that evade MxA restriction, have provided indirect evidence for a subset of the interacting residues at the MxA-NP interface. Evolutionary analysis of MxA orthologs across 24 primate species revealed a clustering of positively selected residues in the stalk region of the protein, specifically the L4 loop, which confer species-specific antiviral activity^44^. This high density of rapidly evolving residues in L4 suggested that this protein region may engage with evolving viruses. Indeed, among the five positively selected L4 residues (G540, F561, F564, S566, and S567), an aromatic residue at position 561 was shown to be especially critical for MxA restriction of H5N1 IAV and THOV^44^ by dictating binding specificity to their NPs. W561 enhanced restriction of H5N1 IAV but appeared to prohibit THOV restriction, whereas Y561 enhanced restriction of THOV^43,45^, demonstrating a breadth-specificity tradeoff in which an optimal MxA sequence against one virus is sub-optimal for another.

Conversely, specific residues within THOV and IAV NPs appear to mediate viral sensitivity to MxA restriction. Just two mutations in the JOSV NP relative to the THOV NP (G327R and R328V) are sufficient to mediate viral escape from MxA restriction both *in vitro* and *in vivo*^7^. MxA escape by 2009 H1N1 (compared to H5N1) required the combined effect of ten mutations in its NP^46^, whereas just three mutations in the 1918 H1N1 NP were sufficient to escape human MxA restriction^46^. Intriguingly, one of the NP mutations required for MxA escape in 1918 H1N1 (L283P) was also shown by a deep mutational scanning analysis to be necessary for the replication of the H3N2 IAV strain in the presence of human MxA^47^.

Despite NP being the best-characterized viral target of MxA and multiple structural analyses of MxA and other Mx proteins^33,48,49^, we still lack direct biochemical insight into the MxA-NP interface. This is partly because native MxA forms dimers, tetramers, and higher-order structures in solution and associates with intracellular membranes^33,34,50^. X-ray crystallographic analysis required the MxA protein to be engineered with four mutations to prevent oligomerization (YRGR440-443AAAA) and a deletion of nearly the entire unstructured L4 domain^33^, which later studies showed to be a critical determinant of NP binding and restriction by MxA. Previously described structures of THOV and IAV NP^51–53^ revealed that these viral proteins form trimers, assemble into helices bound to viral RNA^53,54^, and associate with the viral polymerase complex^51^, adding an additional layer of complexity. For these reasons, MxA-NP modeling efforts have been unsuccessful. As a result, details of the MxA-NP binding interfaces remain mostly unknown.

To address this critical gap in our knowledge, here we performed molecular modeling of the MxA L4 loop in complex with THOV NP. We chose to focus on the THOV NP-MxA interface because THOV is the orthomyxovirus most potently restricted by human MxA. Since previous studies have shown that binding affinity correlates with restriction potency^43^, we reasoned that human MxA may have the highest binding affinity for THOV NP and provide the best opportunity to model the MxA-NP interaction. Using AlphaFold3, we generated a co-folded model of this complex and evaluated its stability through classical all-atom molecular dynamics simulations. We extended this MxA-THOV NP monomer model to potential MxA binding to an NP oligomer. We further modeled key variants at residue 561 and found that our model accurately predicts their relative binding stability to THOV NP. To further assess our model’s predictive accuracy, we identified several MxA mutations predicted to increase MxA restriction, as well as NP mutations predicted to increase viral escape from MxA restriction. We validated these predictions using a minireplicon assay that demonstrated NP variant stability and escape from MxA, as well as increased restriction by MxA variants. Together, our combined modeling and experimental studies provide critical insight into how MxA L4 variants bind an important viral target, how this virus might have evaded MxA restriction in the past, and how it could evolve to evade MxA in the future.

## Results

### The human MxA L4 Loop adopts an unstructured conformation

Human MxA has a tripartite structure comprising an N-terminal GTPase module, a central bundle signaling element (BSE), and a distal stalk domain, which includes a protruding specificity-determining loop L4 that recognizes viral targets (Fig. S1A-B). Despite considerable interest in Mx biology and structure, the MxA L4 loop has not been characterized structurally due to its deletion from MxA to prevent oligomerization in the resolved crystal structures^33,48,49^. Based on evidence of recurrent positive selection and functional importance in viral target binding^43–45^, we hypothesized that the L4 loop of MxA is either unstructured or highly flexible, particularly near the critical residue 561. Such flexibility would be consistent with Loop L4’s dual role as a plastic interface for interaction with different viral targets^55^, and as a membrane-binding domain analogous to the pleckstrin homology domain of dynamin^56^.

However, there is no direct experimental evidence showing that the L4 loop is partially or fully disordered. Instead, AlphaFold3 (AF3) modeling of full-length MxA and full-length MxA/NP complexes consistently predicts that the MxA L4 loop adopts a highly structured conformation, forming a long α-helix spanning residues 533-557 (Fig. 1A), which is also consistent with secondary-structure predictions (Fig. S1C). However, we were cautious about this interpretation because AlphaFold models can be highly biased toward well-structured domains and may overpredict secondary structure in intrinsically disordered or flexible regions^57^. Moreover, in this instance, AlphaFold models could be heavily influenced by the α-helical structure of the L4 loop of the MxB paralog^58^, which has a completely different antiviral range.

**Figure 1.**
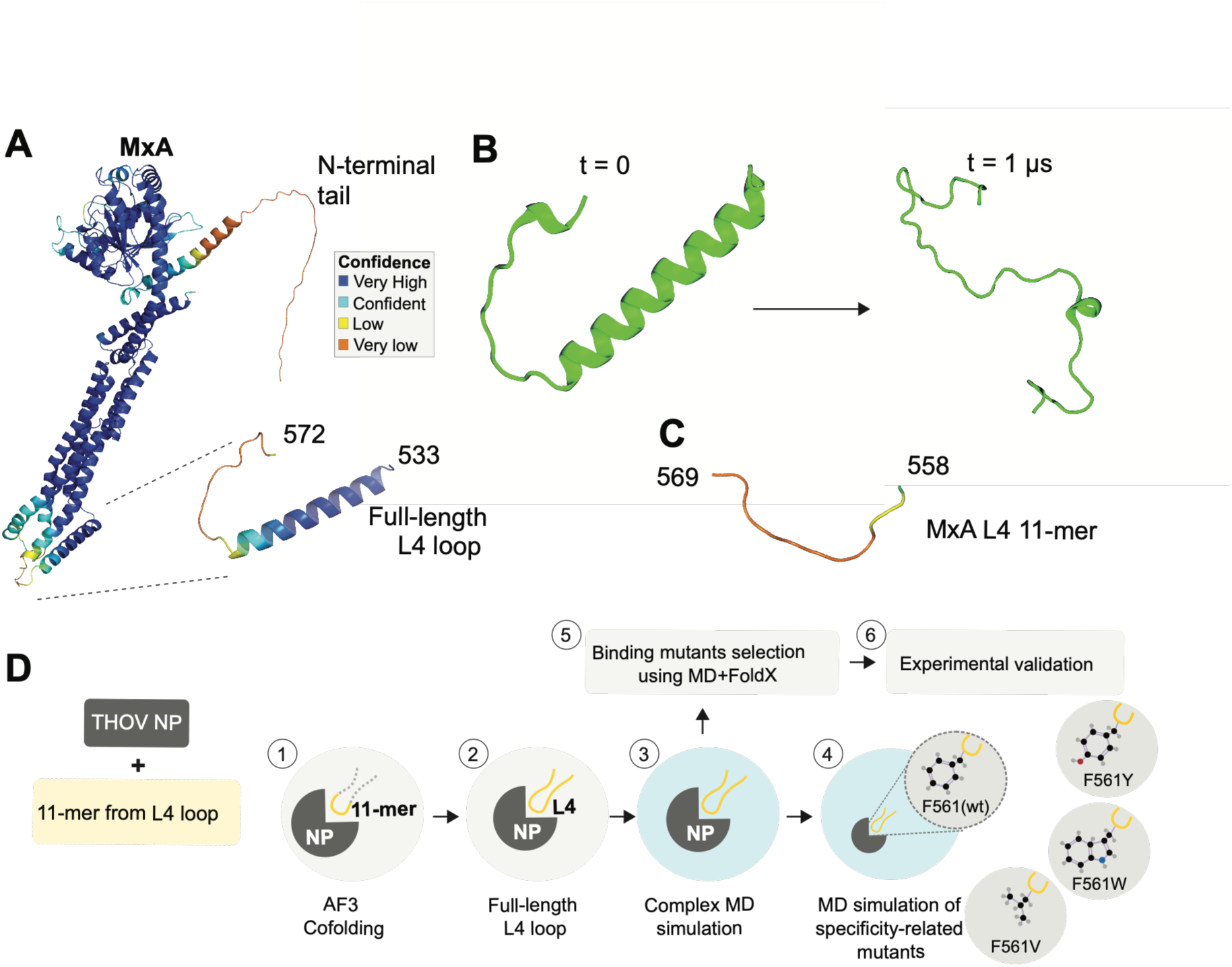
Molecular Dynamics (MD) simulations predict an unstructured human MxA Loop L4, contrary to AlphaFold3 predictions. **(A)** Top-ranked AF3 model of full-length MxA colored by the predicted Local Distance Difference Test (pLDDT), with a close-up view of the L4 region (residues 533–572); colors indicate AF3 confidence levels. **(B)** L4 loop structure at the beginning of the MD simulation (t = 0) and after 1 μs, showing the rapid loss of α-helical secondary structure. **(C)** AF3 model of only the L4 loop 11-mer peptide (residues 558–569); colors indicate AF3 confidence metrics. **(D)** Multi-step pipeline for the computational identification and experimental validation of residues involved in MxA L4-THOV NP recognition. (Step 1) AlphaFold3 co-folding of the THOV NP monomer with an 11-mer peptide derived from the MxA L4 loop to identify an initial binding pose. (Step 2) Construction of a full-length L4 loop model while preserving the binding interface predicted by the top-ranked 11-mer AF3 complex. (Step 3) Molecular dynamics simulations of the THOV NP-L4 loop complex to evaluate the stability of the predicted interaction and characterize the binding interface. (Step 4) Molecular dynamics simulations of specificity-related residue 561 variants (F561V, F561W, and F561Y) to assess their effects on binding. (Step 5) Selection of candidate mutations using predicted binding stability calculations using MD+FoldX. (Step 6) Experimental validation of computationally predicted mutations affecting THOV NP recognition.

To test these preliminary AF3 predictions, we carried out a 1µs-long molecular dynamics (MD) simulation of the L4 loop (MxA residues 533-572); this computational technique models atomic motions over time to assess structural stability and conformational changes. The simulation indicated that, even if the initial AF3 prediction of a helical L4 loop were correct, the helix gradually unfolded, ultimately adopting a flexible, disordered conformation (Fig. 1B). Thus, the MD simulation supported the hypothesis that the MxA L4 loop is disordered and highly flexible, at odds with the AF3 and secondary structure predictions.

Given this prediction disparity, we modeled a shorter 11-residue segment (residues 558-569) of MxA L4, hereafter referred to as the MxA L4 11-mer. The MxA L4 11-mer also includes four of the five human MxA residues previously identified as evolving under strong positive selection in primate evolution: F561, F564, S566, and S567^44^. This segment was consistently predicted as disordered by all three predictions: AF3, a 1µs long MD simulation, and a secondary structure predictor (Fig. 1C and Fig S1). To gain molecular insight into the MxA-NP interaction, we followed a multi-step pipeline consisting of both computational predictions and experimental validation starting with THOV NP interaction with the MxA L4 11-mer (Fig. 1D).

### MxA L4 11-mer appears to interact with THOV NP at a surface distinct from its RNA-binding groove and oligomerization surface

We built structural models of the human MxA L4 11-mer in complex with THOV NP (Step 1, Fig. 1D) using AF3^59^ co-folding of THOV NP with residues 558-569 of the MxA L4 loop. The top-ranked model, selected based on interface-pLDDT, ipTM, and pTM scores, exhibited moderate confidence at the predicted binding interface (Fig. 2A-B). Despite local uncertainty at the interface, the AF3-predicted THOV NP structure closely matched the experimentally determined crystal structure^53^ (RMSD < 1 Å; Fig. S2), supporting the model’s overall structural accuracy. To preserve the co-folded interface with the MxA L4 11-mer, we focused on the AF3-predicted NP structure for subsequent analyses. Based on this AF3 model, we identified viral NP residues predicted to be in contact with the MxA 11-mer to define the MxA-binding interface on THOV NP, which includes surface-exposed residues consisting of small hydrophobic patches surrounded by polar and charged residues that cluster within five structural segments of the NP body (Fig. 2C, S3). Notably, the predicted MxA-binding interface overlaps two critical sites, G327 and R328, which were previously shown to mediate JOSV escape from MxA restriction^7^, despite prior knowledge of these sites not being incorporated into our AF3 modeling.

**Figure 2.**
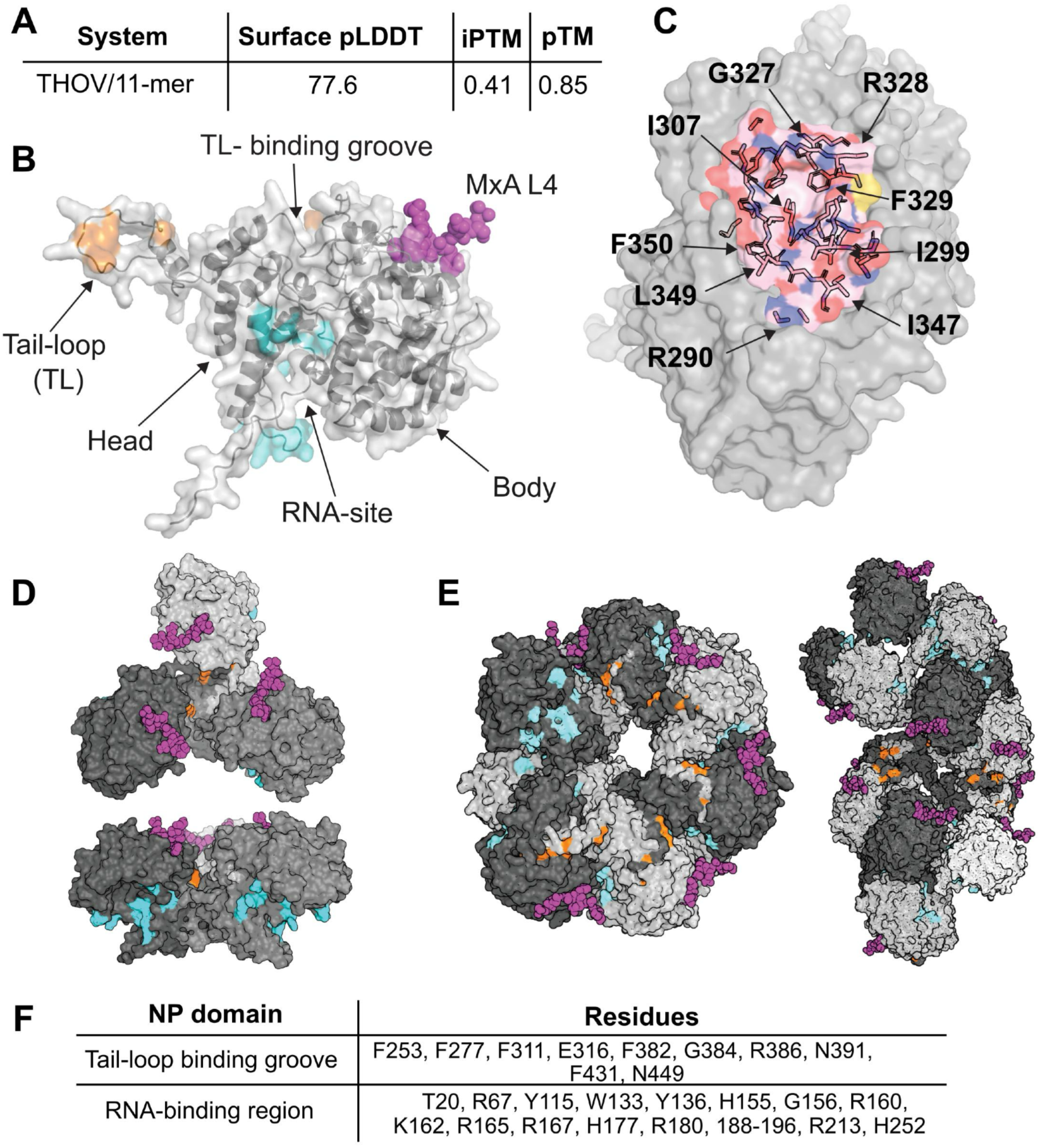
Structural characterization of the THOV NP–MxA interaction and mapping of functional regions. **(A)** AlphaFold3 confidence metrics for the THOV NP monomer in complex with the MxA L4 11-mer complex are presented, including interface pLDDT, ipTM, and pTM scores. **(B)** Structure of a THOV NP monomer showing the major structural regions: head, body, RNA-binding site (cyan), Tail-loop, and TL binding groove (orange). The predicted bound MxA 11-mer peptide is shown in magenta spheres. **(C)** Close-up view of the predicted 11-mer binding site on THOV NP. Residues predicted to interact with the MxA 11-mer are labeled and colored by physicochemical properties: positively charged (blue), negatively charged (red), and hydrophobic (pink). **(D)** Two orientations of the THOV NP trimer are shown (PDB: 8CJW)^53^. THOV NP oligomerization occurs by the NP monomer tail loop inserting into a groove (shown in orange) in the neighboring NP monomer^53^. THOV NP RNA-binding residues are shown in cyan, and the MxA 11-mer is shown by magenta spheres. **(E)** THOV NP ribonucleoprotein assembly^53^ showing the spatial distribution of the tail-loop binding groove (orange), RNA-binding residues (cyan), and the predicted MxA 11-mer (magenta spheres): RNA is not shown. The predicted MxA-binding site is exposed on the surface of the oligomeric assembly. It is distinct from and does not occlude the canonical RNA-binding and oligomerization regions. **(F)** We tabulate the residues^45^ that make up the tail-loop binding groove and RNA-binding regions on THOV NP. These residues do not overlap with the predicted MxA L4 binding site on THOV NP, highlighted in (B).

Previous studies have proposed that MxA restricts orthomyxoviruses by forming higher-order oligomers that encircle viral nucleocapsids, sterically blocking essential interactions within the vRNP complex^32,33^. We therefore asked whether the MxA-binding site predicted by our model would still remain accessible in oligomeric NP assemblies. To address this question, we modeled the MxA L4 11-mer complex with trimeric or higher-order THOV NP assemblies from previously obtained experimental structures^53^ using restraint-guided docking. Our analysis shows that the MxA 11-mer binding surface is spatially distinct from the surfaces involved in either RNA-binding or NP oligomerization interfaces (Fig. 2D-E). In THOV NP, the RNA-binding groove^45^ forms a continuous concave surface between the NP head and body domains, whereas oligomerization occurs through the tail-loop binding groove^45^ (Fig. 2B and 2F). In contrast, the predicted MxA-binding site on THOV NP (including residues L283 and Y289) is located on the outer surface of the head domain, where it remains accessible to the full-length MxA protein. This interface location would allow MxA oligomers to dock onto higher-order NP assemblies without directly occluding either the RNA-binding or oligomerization surfaces. Our findings provide a direct model of how the MxA loop L4 might engage with the THOV NP at the molecular level (Fig. 2C-E).

### Molecular modeling of full-length MxA L4 interaction with THOV NP confirms several aspects of MxA restriction of THOV

To determine whether the interactions identified for the MxA L4 11-mer are preserved in the context of the complete loop, we next modeled the full-length MxA L4 loop (residues 533–572) in complex with THOV NP using homology modeling based on the top-ranked AF3 11-mer model (Step 2, Fig. 1D; Fig. S4). The resulting complex of THOV NP with the unstructured full-length L4 loop was subsequently refined and evaluated by MD simulations (Step 3, Fig. 1D; Methods). MD simulations showed that extending the L4 loop did not disrupt the interactions formed by the central 11-mer. Instead, the 11-mer remained stably associated with THOV NP throughout the simulation, as indicated by low RMSD values and persistent positioning of residue F561 within the predicted NP-binding pocket (Fig. S5).

Consistent with these observations, per-residue root mean square fluctuation (RMSF) analysis showed minimal flexibility around residues 558-561 (Fig. 3A). More broadly, residues 554– 564 surrounding F561 maintained close contacts with THOV NP, exhibiting small mean inter-residue distances (Fig. 3B) and moderate-to-high contact frequencies throughout the simulation (Fig. 3C). In contrast, the remaining L4 loop residues exhibited greater conformational flexibility, particularly at the loop termini, which formed only transient contacts with THOV NP and showed a low contact frequency. One notable exception was at residues close to the L4 loop N-terminus, which exhibited relatively high contact frequencies for residues 537-539 and a moderate contact frequency for residue 540, comparable to that of residue 561 despite their greater distance from the central binding motif. This is intriguing because residue 540 is one of five residues in the MxA L4 loop identified as undergoing recurrent positive selection during primate evolution^44^ and was previously shown to be critical for MxA interaction with THOV NP^43^. Its apparent contact with THOV NP in our simulation is consistent with these previous findings.

**Figure 3.**
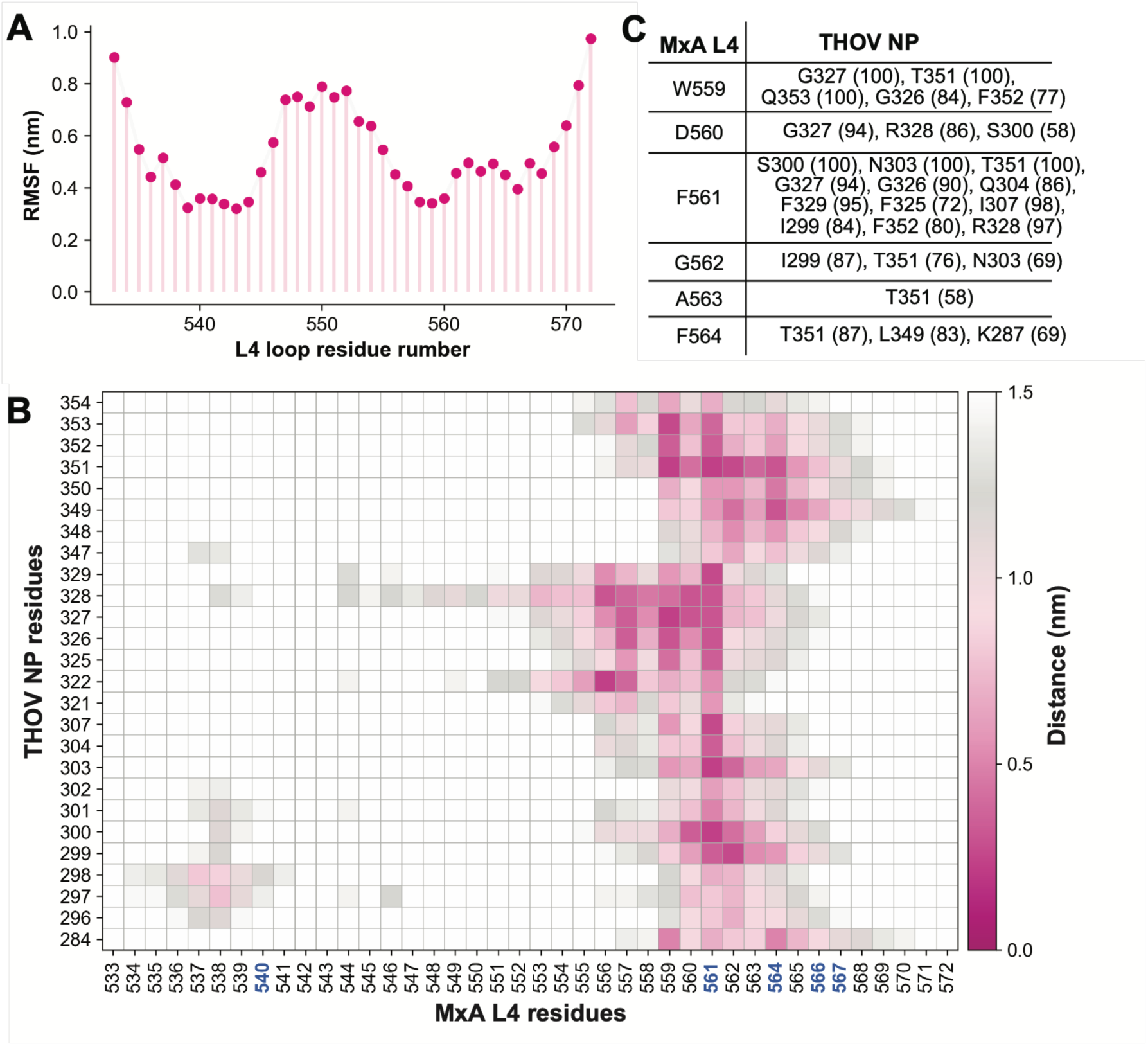
Dynamic and structural characterization of the MxA L4 loop interaction with THOV NP. **(A)** Root mean square fluctuation (RMSF) profile (fluctuations around the mean position over time) of the MxA L4 loop (residues 533–572) during a 200 ns MD simulation of the complex. **(B)** Distance heatmap showing the average minimum distance between THOV NP residues and MxA L4 residues throughout the simulation (time-averaged pairwise distances), ranging from 0 (tight contact) to > 1.5 nm (no measurable contact); darker colors indicate closer contacts. MxA L4 residues that have evolved under positive selection during primate evolution are highlighted on the X-axis (blue). THOV NP residues are indicated on the Y-axis. **(C)** Summary of high-occupancy contacts between MxA L4 loop residues and THOV NP. Values in parentheses indicate the percentage of simulation frames in which each residue pair remained in contact during the MD simulation (higher numbers indicate tighter contact).

We next examined specific residue-level pairings critical for MxA L4 loop interaction with THOV NP. Analysis of the MD trajectories revealed several L4 loop residues, especially residues 559 through 564, which consistently stayed proximal to viral NP residues (Fig. 3B). Among these, the MxA residue F561, previously identified as critical for orthomyxovirus restriction, serves as a central anchoring point, forming high-occupancy contacts with THOV NP residues S300, N303, I307, G327, R328, F329, and T351 (Fig. 3B-C). Many of the contacted NP residues maintained >80% contact frequency throughout the simulations. Notably, a few of these NP residues, such as G327 and R328, correspond precisely to positions that have been experimentally associated with MxA escape or sensitivity^7^, even though these constraints were not built into our model *a priori*. NP residue G327 has a high contact frequency with MxA W559, D560, and F561, whereas NP residue R328 has a high contact frequency with MxA D560 and F561 (Fig. 3C). Overall, these residue-level interactions provide the first detailed models of how critical MxA L4 loop residues interact with the THOV NP surface.

Among all Loop L4 residues, the phenylalanine residue at position 561 in wild-type human MxA makes the dominant contribution to THOV restriction^43,44,55^. Previous mutagenesis studies showed that only aromatic hydrophobic residues (F, Y, or W) at this position retain antiviral activity, with a clear functional hierarchy in restriction potency (Y561 > F561 > W561), whereas substitution with any other amino acid abolishes restriction^43,45^. Because our structural models identified MxA residue 561 as a key anchoring residue at the MxA-THOV NP interface, we next asked whether MD simulations and interaction energy analyses could recapitulate this functional hierarchy (Step 4, Fig. 1D). We therefore performed MD simulations of four variants at position 561: the three restriction-competent residues (Y, F, and W) and the restriction-deficient variant V. Consistent with experimental observations, we found that both the F561Y and F561 variants maintained stable proximity to the THOV binding pocket throughout the simulation (Fig. 4A), whereas F561W initially had robust interactions but ultimately lost contact and dissociated from the site. In contrast, F561V failed to maintain contact and detached early in the simulation. These results indicate that tyrosine substitution at position 561 stabilizes the interaction with THOV NP more effectively than either the wild-type phenylalanine or the tryptophan variant, likely due to improved hydrogen bonding and electrostatic complementarity (Fig. 4B-C). This conclusion was supported by energy decomposition analyses (Fig. 4D), which showed that F561Y significantly enhanced both van der Waals (-70.0 ± 0.17 kJ/mol, Lennard-Jones potential energy) and electrostatic interactions (-61.1 ± 0.26 kJ/mol, Coulomb interaction enegy), resulting in the most favorable total interaction energy (-131.1 ± 0.31 kJ/mol) among the three ‘active’ variants. These findings are consistent with previous combinatorial mutagenesis studies, which have shown that F561Y substitutions enhance THOV restriction, whereas F561W substitutions are, on average, incompatible with THOV restriction^43,45^.

**Figure 4.**
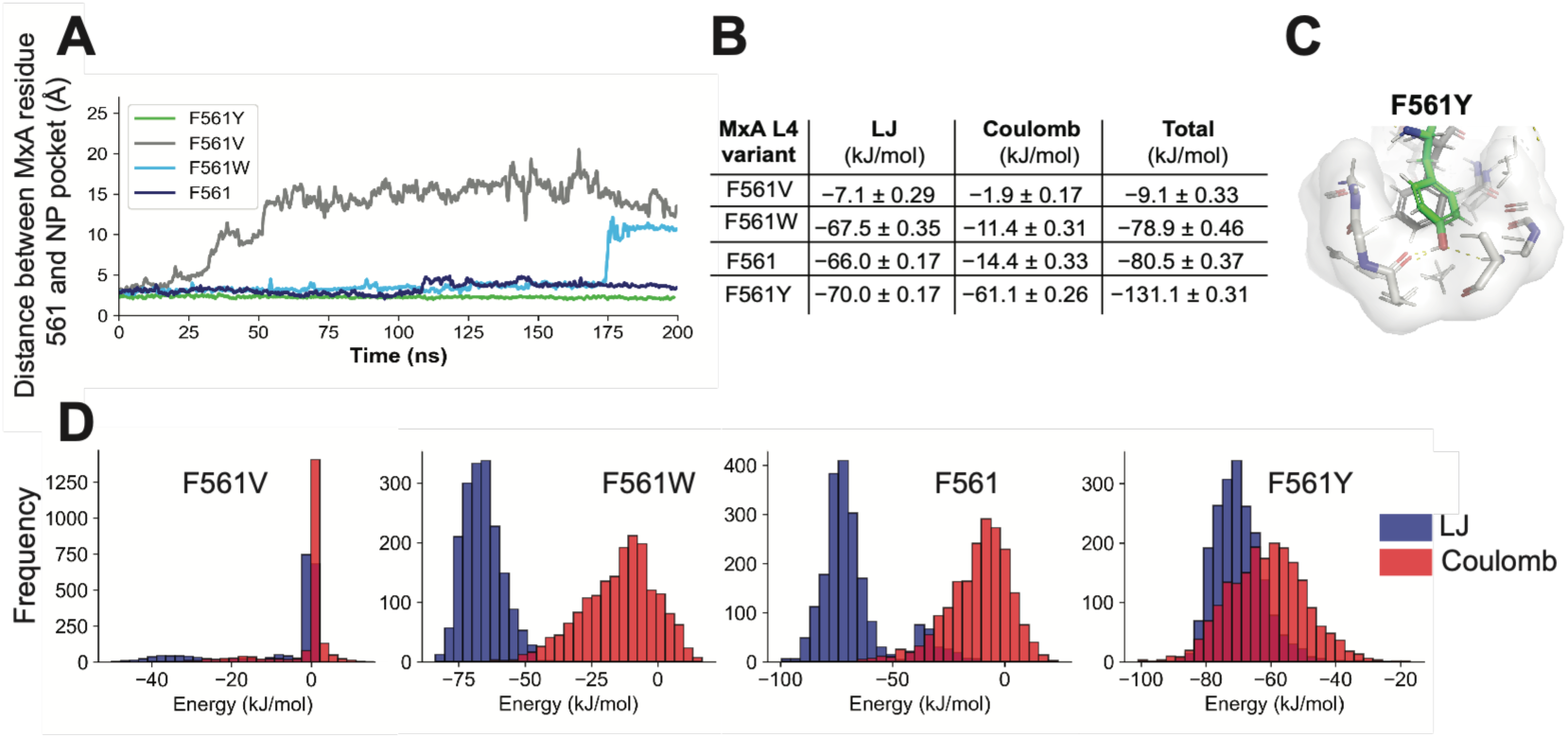
Modeling the effects of residue 561 substitutions on THOV NP-MxA L4 loop interactions. **(A)** Distance between residue 561 in the L4 loop and the NP binding pocket during 200 ns MD simulations for the wild-type residue (F561) and the F561V, F561W, and F561Y variants. F561Y displays the strongest overall interaction energy, followed by wildtype F561, then F561W, which binds stably until 175 ns, after which it loses binding. F561V has the lowest overall interaction. **(B, D)** Energy decomposition analysis of residue 561 interactions with the binding pocket for the F561V, F561W, wild-type F561, and F561Y variants. Histograms show the distributions of Lennard-Jones (van der Waals, blue) and Coulombic (electrostatic, red) interaction energies sampled throughout the simulations. **(C)** The F561Y variant displays the strongest overall interaction energy due to additional electrostatic stabilization. This enhancement is consistent with the formation of hydrogen-bonding interactions and improved electrostatic complementarity between the tyrosine hydroxyl group and residues within the binding pocket.

### *In vitro* MxA restriction of THOV NP corroborates computationally predicted stabilizing and destabilizing mutations at the MxA L4 -THOV NP binding interface

Based on our structural models of the MxA L4 loop in complex with THOV NP, we hypothesized that *in silico* mutational analysis of the MxA L4 loop-NP interaction interface could help identify candidate mutations that enhance or impair the stability of the MxA L4 loop-NP binding interface, thereby affecting the potency of the MxA restriction of THOV. To do this, we first performed comprehensive *in silico* mutational scans of all possible variants in each MxA loop L4 residue (533-572) (Step 5, Fig. 1D, Dataset S1). Since we do not have a pooled assay to measure restriction of all these MxA variants, we selected five representative mutations to test in a minireplicon assay (below). We prioritized mutations with favorable predicted relative binding free energy values (ΔΔ*G*_bind_ < 0) and high contact occupancy during MD simulations. We also selected variants in different positions of the L4 loop, intentionally selecting positions that had not been previously tested in antiviral assays, *e.g.,* avoiding the five positively selected L4 positions that had previously been tested in combinatorial mutagenesis screens^43,44^. Finally, we tested variants that differed from the cognate residue in wild-type MxA by diverse physicochemical properties. From these analyses, we selected five MxA missense variants with high potential to improve binding to THOV NP for testing: V537L, K557W, S568R, T570P, and S572E (Fig. 5A). These residues are highly conserved among primate MxA orthologs (Fig. S6A-B).

**Figure 5.**
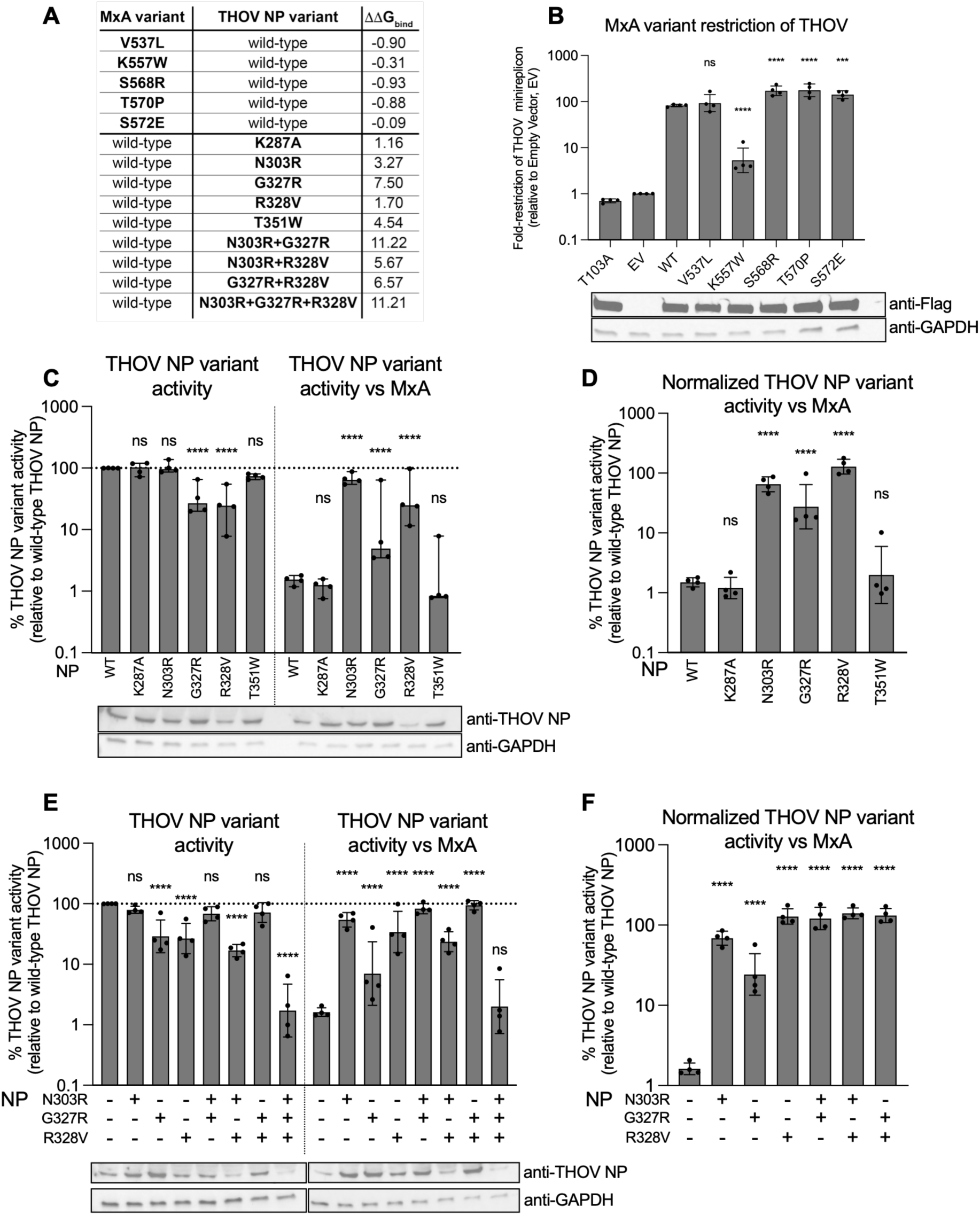
Testing MxA-NP binding predictions using minireplicon restriction assays. **(A)** Selected mutations predicted to improve (negative ΔΔG_bind_) or worsen binding of THOV NP (positive ΔΔG_bind_) are shown. **(B)** Fold restriction of different MxA variants relative to empty vector (EV) using a minireplicon luciferase assay. Empty Vector (EV), a catalytically dead GTPase MxA (T103A), and wild-type (WT) MxA are shown as controls, and asterisks indicate statistical significance of the difference between fold restriction of the THOV minireplicon by variants and that mediated by WT MxA. Western blots indicate expression levels of different flag-tagged MxA variants relative to a GAPDH loading control. **(C)** Activity of THOV NP variants in the minireplicon assay without MxA (left) and in the presence of MxA (right), relative to wild-type (WT) THOV NP. Western blots show expression levels of different THOV NP variants relative to a GAPDH loading control. **(D)** Normalized activity of the different THOV NP variants in the presence of MxA relative to activity in the absence of MxA (C). **(E)** Activity of single-, double-, and triple-mutant THOV NP variants, relative to wild-type (WT) THOV NP, in the absence of MxA (left) or in the presence of MxA (right). Western blots show expression levels of different THOV NP variants relative to a GAPDH loading control. **(F)** Normalized activity of the single-, double-, and triple-mutant THOV NP variants in the presence of MxA relative to activity in the absence of MxA (E). Four replicates were performed for each mutant. All data were log-transformed prior to two-way ANOVA without an interaction term, with Dunnett’s post-hoc test. Individual data points represent independent replicates, and error bars indicating the geometric mean and geometric SD are shown. Asterisks indicate statistical difference compared to the mean fold-restriction by WT MxA or compared to the activity of wild-type THOV NP in the same conditions. *p<0.05, **p<0.01, ***p<0.001, ****p<0.0001.

Although the MxA variants we selected were predicted to improve binding to THOV NP, co-immunoprecipitation assays to directly measure differences in binding affinity between MxA variants and THOV have a relatively narrow dynamic range^43^. Therefore, instead of co-immunoprecipitation assays, we tested the impact of these MxA mutations using a THOV minireplicon assay, which has a much broader dynamic range. This assay measures NP-dependent viral polymerase activity without requiring infectious virus (Step 6, Fig. 1D; Methods)^60^. We quantified restriction by each MxA variant against THOV relative to both an empty vector control and a restriction-deficient MxA mutant (T103A)^61^ and compared this activity to that of wild-type MxA, which already potently restricts THOV^62,63^. Western blot analyses confirmed that all five MxA variants are expressed at levels that are comparable to wild-type MxA and the T103A catalytically dead variant. Thus, any changes in activity are not the result of altered expression or stability. Of the five MxA variants selected for experimental validation, one (V537L) had no discernible effect. This may be because it represents a conservative hydrophobic substitution in which the small increase in side-chain size does not substantially alter the local interaction. One selected variant (K557W) reduced antiviral activity relative to wild-type MxA, potentially because replacing K557 with a bulky hydrophobic tryptophan disrupts the electrostatic interaction with the nearby THOV NP residue E322 (Fig. 5B and Fig. 3B). However, three of five candidate mutations prioritized solely on the basis of our predicted MxA-THOV NP interaction model (S568R, T570P, and S572E) significantly enhanced MxA restriction of THOV, supporting the predictive power of our structural model.

We applied the same computational prioritization strategy as with MxA L4 loop mutations to identify potential viral escape mutations in THOV NP. For this, we focused on 62 NP residues located within 10 Å of the predicted MxA L4 binding site to identify variants predicted to weaken binding (positive ΔΔ*G*_bind_ values) to human MxA (Dataset S2). The vast majority of variants either did not significantly improve predicted binding affinity (negative or neutral ΔΔ*G*_bind_ values) or were predicted to be strongly misfolded (high ΔΔ*G*_fold_). From the remaining variants, we selected five missense variants in THOV NP with a range of positive ΔΔ*G*_bind_ values predicted to enable viral escape from restriction: K287A, N303R, G327R, R328V, and T351W (Fig. 5A). These residues are variable across NP proteins from THOV and THOV-related viruses (Fig. S6C-D). Two of these mutations have already been previously associated with escape from human MxA restriction in the JOSV virus (G327R, R328V)^7^, whereas the other three represent previously untested candidates.

Because the minireplicon assay relies on measuring NP-dependent polymerase activity, we first determined whether any of these five mutations affected baseline NP function even in the absence of MxA. For this, we used the minireplicon assay to assess NP-dependent polymerase activity in the absence of MxA (using an empty vector control, EV). Western blot analyses revealed no gross alterations in NP expression or stability for most variants, with the possible exception of R328V. We found that three of five THOV NP variants tested (K287A, N303R, and T351W) had no discernible change in NP-dependent polymerase activity in the absence of MxA (Fig. 5C), indicating they had no adverse effects on NP activity. In contrast, previously described naturally occurring escape variants G327R and R328V in JOSV each resulted in ∼5-fold reduction in NP-dependent polymerase activity, suggesting that these single missense mutations may incur a fitness cost to the virus (Fig. 5C). Both substitutions were also predicted to destabilize NP folding based on their positive Δ*ΔG*_fold_ values (7.5 kcal/mol for G327R and 1.8 kcal/mol for R328V), consistent with their reduced polymerase activity.

We next tested whether any of these NP variants could escape restriction by wild-type MxA in the minireplicon assay. We found that two variants, K287A and T351W, did not substantially differ from wild-type THOV NP in their susceptibility to MxA restriction. However, the remaining three variants (N303R, G327R, and R328V) all conferred escape (Fig. 5C). By normalizing their NP-dependent polymerase activity in the presence versus absence of MxA restriction, we show that N303R, G327R, and R328V encode 43-fold, 18-fold, and 85-fold escape from MxA restriction, respectively, relative to wild-type THOV NP (Fig. 5D). Overall, three out of five computationally predicted NP variants conferred escape from MxA restriction in an *in vitro* setting, providing strong experimental support for the predictive power of our structural model. Importantly, the N303R mutant is entirely novel and mediates an escape as strong as that of the naturally occurring JOSV escape variants G327R and R328V but without incurring fitness costs discernible in the minireplicon assay, demonstrating that our computational framework can identify novel viral escape mutations.

Viral escape from MxA often involves fitness-lowering mutations that can be subsequently compensated for by secondary mutations^7,46^. We used MD+FoldX to predict the impact of the possible combinations of the N303R, G327R, and R328V mutations that most significantly led to MxA escape (Fig. 5A). This MD+FoldX analysis predicted that some mutant combinations would further weaken MxA binding, thereby enhancing viral escape (Fig. 5E). Using minireplicon assays, we found that two double mutant combinations (N303R/ G327R and G327R/ R328V) restored full baseline NP activity, whereas the baseline activity of the N303R/R328V double mutant remained as functionally impaired as the R328V single mutant (Fig. 5E). This suggests that positive epistasis between N303R and G327R, or between G327R and R328V (the combination found in JOSV) can restore full NP functionality in addition to mediating MxA escape. In contrast, the triple mutant (N303R/G327R/R328V) was severely impaired in baseline NP activity (Fig. 5E), consistent with its ΔΔG_fold_ value of 4.8 kcal/mol and reduced expression or stability evidenced by western blot. We found that all NP double mutants also conferred a high degree of MxA escape in minireplicon assays (Fig. 5E). Normalizing to their respective baseline NP activities, we infer that all double mutants encode ∼80-fold higher escape than wild-type THOV NP (Fig. 5F). Since the baseline NP activity is so impaired in the triple mutant, we cannot reliably infer the extent of its escape from MxA restriction.

Together, our MD-guided minireplicon assays identify three novel MxA variants that improve restriction, validate two previously identified NP escape variants, and identify one novel THOV NP variant that escapes MxA restriction. This supports the predictive potential of our NP-MxA loop L4 complex model to identify mutations that enhance THOV NP escape from MxA or enhance MxA binding to THOV NP and thereby enhance restriction.

## Discussion

Emerging orthomyxoviruses continue to test the limits of mammalian innate immunity, placing the interferon-induced GTPase MxA at the center of a long-standing evolutionary arms race. Although influenza A virus (IAV) strains have rightly attracted most attention, JOSV exemplifies how just two mutations in a tick-borne orthomyxovirus are sufficient to bypass the MxA restriction factor and potentially breach the species barrier to cause zoonosis^7^. Yet, despite decades of work on Mx proteins and viral nucleoproteins (NPs), the molecular architecture of the MxA-NP interface has remained stubbornly opaque. Our study addresses this gap by providing the first structural insight into how the rapidly evolving MxA L4 loop engages THOV NP and how subtle changes on either side of this interface rewire the outcome of host-virus conflict.

One of the most notable findings of this study is that the MxA L4 loop is unlikely to adopt the stable α-helical conformation initially predicted by AlphaFold3. Instead, MD simulations consistently favored a flexible, intrinsically disordered conformation that better reconciles the rapid evolution, positive selection, and broad target specificity of this region (Fig. 1). By deliberately restricting the modeled segment to an 11-residue L4 “micro-interface,” we captured a compact, dynamic binding mode that is compatible with both positive selection patterns in primate MxA and prior mutational data. This framework offers a general strategy for interrogating intrinsically flexible antiviral loops, which frequently mediate host-virus interactions but remain challenging to resolve using current structure prediction methods alone (Fig. 1D).

Our structural model identifies a previously unrecognized MxA-binding surface on THOV NP that is spatially distinct from both the RNA-binding groove and the oligomerization interface (Fig. 2) and is unlikely to affect either of these properties. Instead, MxA binding may interfere with the higher-order organization, transport, or functional dynamics of viral ribonucleoprotein complexes required for productive replication. Because this interface remains exposed on assembled NP oligomers, our model is also compatible with previously proposed dynamin-like mechanisms^64,65^ in which oligomeric MxA decorates viral nucleocapsids before exerting its antiviral activity. Thus, our findings provide a structural rationale that unifies previously proposed models of MxA restriction with residue-level interactions between the MxA L4 loop and THOV NP.

Previous studies have highlighted the important role of residue 561 in MxA L4 for orthomyxovirus restriction. Our present study confirms residue 561 as a structurally anchored driver of interaction energetics. MD and interaction energy analyses show that F561 serves as a central anchor for MxA L4 onto the NP surface and maintains high-occupancy contacts with multiple NP residues, including at residue sites where mutations can lead to viral escape (Fig. 3). By systematically comparing aromatic and non-aromatic substitutions at 561, we recapitulate the experimentally observed hierarchy of Y561 over F561 and the relative inability of W561 to mediate THOV restriction^43,45^ (Fig. 4). Our study provides a physical explanation: Y561 optimizes both van der Waals and electrostatic complementarity, whereas W561, despite being aromatic, progressively destabilizes the binding interface during simulation. These results showcase how a single positively selected residue can simultaneously tune breadth, potency, and vulnerability to viral counter-adaptation.

Our simulations also identified interactions involving residues near MxA position 540, although these observations should be interpreted with caution because this region lies within a highly flexible portion of the L4 loop that is less well-constrained structurally than the central 11-residue binding motif. Consequently, transient contacts formed by the N-terminal region of L4 may reflect conformational sampling bias rather than persistent interactions within the intact MxA oligomer. In contrast, the interaction network centered on residue 561 is supported by both a more confidently modeled structural region and extensive experimental evidence, making its involvement in THOV NP interaction considerably more robust. Nevertheless, the apparent involvement of residue 540 remains biologically intriguing, as this residue has previously been implicated in THOV NP recognition^43^ and has undergone recurrent positive selection during primate evolution^44^. Future structural and functional studies of full-length MxA will be required to determine whether this region contributes directly to NP recognition or modulates the dynamics of the L4 loop during antiviral restriction.

A particularly compelling aspect of our study is the structural model’s predictive performance. Without incorporating any prior experimental constraints, the predicted MxA L4 interface converged on THOV NP residues G327 and R328, whose mutation enables JOSV to escape MxA restriction *in vitro* and *in vivo*^7^. Building on this structural framework, our MD+FoldX-guided mutational scan workflow prospectively identified both gain-of-host restriction variants in MxA and viral loss-of-binding escape variants in THOV NP (Fig. 5A). Experimental validation using minireplicon tests strongly supported most of these predictions. On the host side, we identified three previously uncharacterized MxA variants (S568R, T570P, S572E) that enhance THOV restriction, demonstrating that even L4 loop residues that are deeply conserved in primate evolution harbor evolutionary potential to increase antiviral potency. On the viral side, we validated both previously known escape mutations from JOSV (G327R, R328V) and uncovered N303R as a novel escape substitution that maintains NP polymerase activity while conferring strong resistance to MxA. The fact that three out of five NP candidates and three out of five MxA candidates behave as predicted indicates that our structural model is sufficiently accurate to guide focused sequence surveillance and rational engineering of MxA variants with increased restriction of THOV. Further optimization of prediction models, for example, by considering full MxA protein folding and complex trafficking, could further improve the predictive power of this workflow. This convergence among unbiased structural prediction and functional validation of restriction and escape variants provides strong orthogonal support for our correct identification of the MxA-THOV NP interaction. More broadly, our findings demonstrate that integrating co-folding with molecular dynamics can both retrospectively explain historical viral escape mutations and prospectively identify new escape variants, providing a framework for engineering more potent MxA proteins to deploy as potential therapeutics while improving surveillance and assessment of the zoonotic potential of THOV and related orthomyxoviruses.

Previous studies have shown that some ‘escape’ mutations that allow viruses to evade MxA restriction negatively affect replication of IAV in the absence of MxA^46,66^. For example, the MxA-resistant P283 variant of IAV NP affected NP stability by making it more aggregation-prone, especially at higher, febrile temperatures^67^. The intrinsic fitness cost of such ‘escape’ mutations further confirms that these very likely arose under strong MxA-mediated selection and were subsequently compensated for by other mutations that epistatically restored viral fitness. Our analyses of single and combined NP ‘escape’ mutations also reveal some instances of epistasis in the MxA-virus arms race. We initially found that two escape mutations (*e.g.,* G327R or R328V) incur fitness costs in the absence of MxA. Our findings are slightly discrepant with the original JOSV analysis, which did not find fitness costs associated with the R328V variant^7^. These differences might reflect differences in expression levels of the R328V variant relative to viral polymerase in our minireplicon assay, compared to the earlier study. Nevertheless, we found that these costs can be mitigated by combining both G327R and R328V variants or by other compensatory substitutions such as N303R, thereby restoring polymerase activity while maintaining high-level escape. Conversely, the lower NP activity (and possibly stability) of the N303R/G327R/R328V triple mutant underscores that not all paths through sequence space are accessible. Viruses must navigate a rugged fitness landscape in which only specific combinations of potential ‘escape’ mutations balance baseline fitness with resistance. Given their context-dependence, these paths are likely to be highly virus-specific. Our combined MD+FoldX and minireplicon framework provides a blueprint for mapping such evolutionary landscapes prospectively for other antiviral effectors and viral nucleoproteins. By identifying structurally constrained yet functionally important interaction surfaces, this approach may also facilitate the rational design of peptide- or protein-based antivirals that target conserved nucleoprotein interfaces.

Our work has direct implications for orthomyxovirus surveillance and comparative pathogenesis. The THOV NP surface we define is structurally conserved across *Orthomyxoviridae* (Fig. S7). Thus, the MxA-binding surface may be conserved between THOV and the emerging Bourbon virus, raising the possibility that analogous escape routes may be available, or already exploited, in related tick-borne viruses. In contrast, influenza A escape mutations cluster on partially distinct NP surfaces^7,46^ (Fig. S8), suggesting that MxA may recognize overlapping but non-identical epitopes on different orthomyxoviral NPs, with residue 561 acting as a key “switch” between binding non-identical surfaces in THOV and IAV NPs. Systematically porting our approach to influenza and other MxA-sensitive viruses could help determine whether a unified structural logic underlies MxA’s broad antiviral repertoire, or whether this effector repeatedly “reinvents” distinct binding solutions across viral families.

## Materials and Methods

### Modeling NP-MxA L4 loop complexes

The NP from Thogotovirus (THOV; UniProt ID: P89216) was co-folded with an 11-residue segment of the human MxA L4 loop (residues 558–569; sequence: SWDFGAFQSSS), which includes sites previously identified as being under strong positive selection^44^. Co-folding was performed using the AlphaFold3 (AF3) algorithm^59^, a state-of-the-art deep learning-based protein structure prediction tool that enables complex modeling of interacting proteins and peptides. Five structural models were generated during co-folding. The quality of each model was assessed using multiple metrics: the average predicted Local Distance Difference Test (pLDDT) score of interface residues (Interface-pLDDT)^68^, the inter-protein Template Modeling score (ipTM), and the predicted TM-score (pTM).

To construct the complete L4 loop, the full-length segment (residues 533–572) was modeled onto the top co-folded NP-11-mer complexes using the automodel class in MODELLER^69^. The resulting model, consisting of a complete unstructured L4 loop, was subsequently analyzed to identify residues that comprise the NP-binding interface, defined as those containing at least one atom within 5 Å of any THOV NP atom. These interface residues were then compared with previously reported sites associated with MxA sensitivity or resistance.

### Modeling the MxA binding interface on multimeric THOV NP assemblies

To investigate the MxA binding interface across multimeric THOV NP assemblies, we carried out restraint-guided protein-protein docking using ClusPro 2.0^70^. Experimental structures of THOV NP in its trimeric form^53^ (PDB: 8CJW) and higher-order multimeric assembly^53^ (PDB: 8RYT) were used as receptors, whereas the AlphaFold-predicted MxA 11-mer structure (model 0) was used as the ligand. Attraction restraints were applied between THOV NP residues 327-328 and MxA residue 561 to guide docking toward the interface identified during our AlphaFold3 co-folding run. For visualization purposes, representative docking poses of the MxA 11-mer were selected for two distinct NP assemblies.

### Assessing the stability of MxA L4-THOV NP complexes

The stability of the wild-type THOV NP–L4 loop complex and corresponding mutants was evaluated using molecular dynamics (MD) simulations. The final THOV NP–full-length MxA L4 loop model generated by MODELLER in the previous modeling step was used as the starting structure for simulations of the wild-type complex. Mutant complexes (F561W, F561Y, and F561V) were generated by introducing the mutations into a representative conformation extracted from the 200 ns wild-type MD simulation trajectory. To further assess the intrinsic stability of the L4 loop secondary structure, simulations were also performed on the isolated L4 loop segment encompassing residues 533-572. All systems were prepared for MD simulations using the CHARMM-GUI^71^ interface. Briefly, each complex was solvated in an explicit water box containing ions at physiological concentration, followed by energy minimization and equilibration. Production MD simulations were subsequently performed using GROMACS 2022.5^72^ for 200ns for protein complexes and 1 μs for the isolated L4 loop simulations. Topologies were created using the parameters from the CHARMM36m^73^ force field. Each complex was placed in a rectangular simulation box such that all protein atoms were at least 1 nm from the box edges, and the box was then filled with TIP3P^74^ water molecules. Each box was neutralized by adding the appropriate number of Cl^−^ and Na^+^ counterions at a concentration of 0.15 M. Systems were subsequently energy-minimized under periodic boundary conditions using the steepest-descent algorithm^75^.

An initial equilibration phase of 1 ns was performed under NVT conditions to ensure that water molecules equilibrated around the proteins. The equilibration phase began at 0 K with a linear temperature increase to 303.15 K. A subsequent equilibration phase was performed under NPT conditions by enabling pressure coupling at a reference pressure of 1 bar for 1 ns. Position restraints were applied during equilibration, with force constants of 400 kJ/mol·nm² for backbone heavy atoms and 40 kJ/mol·nm² for side-chain heavy atoms. Production simulations were then conducted in the NPT ensemble at 303.15 K and 1 bar with a 2-fs time step. Periodic boundary conditions were applied in all stages, and hydrogen bonds were constrained using the LINCS^76^ algorithm. Temperature and pressure were controlled using the Nosé-Hoover^77^ thermostat and the Parrinello-Rahman^78^ barostat, respectively, during production simulations. Electrostatic interactions were calculated using the Particle Mesh Ewald^79^ method with a real-space cutoff of 1.2 nm, and van der Waals interactions were treated using the same cutoff. Stability was assessed using energy contribution analysis, hydrogen-bond quantification, Root Mean Square Deviation (RMSD) calculations, and measurements of the distances between key MxA residues and the viral nucleoprotein.

### Selecting mutants for experiments

Building on the MD+FoldX framework established in our previous studies^80,81^, we estimated the effects of mutations at the wild-type NP-L4 loop interface on both folding (ΔΔ*G*_fold_) and binding stability (ΔΔ*G*_bind_). An *in silico* single-site mutational scan was performed using FoldX 5.1^82^ on all residues located at the interaction interface, defined as all L4 loop residues (40 residues × 19 possible amino acid changes) and NP residues within 10 Å of the L4 loop (62 residues × 19 possible amino acid changes). Prior to mutation modeling, the geometry of the wild-type complex was optimized using the *RepairPDB* command, followed by six rounds of energy minimization under the FoldX force field. The binding free energy of the wild-type complex (Δ*G*_wt_) was calculated using the *AnalyseComplex* command. Point mutations were then introduced using the *BuildModel* routine, and the binding energy of each mutant complex (Δ*G*_mut_) was again computed with *AnalyseComplex*, from which ΔΔ*G*_bind_ (= Δ*G*_mut_ - Δ*G*_wt_) was derived. Folding stability (ΔΔ*G*_fold_) was assessed for NP residues only. For both ΔΔ*G*_bind_ and ΔΔ*G*_fold_, we report averages calculated across three representative snapshots from the MD simulations to account for structural variability. Mutations were classified as disruptive if they exhibited high ΔΔ*G*_bind_ and low folding stability (ΔΔ*G*_bind_ > 0.5 kcal/mol and ΔΔ*G*_fold_ < 0.5 kcal/mol), and as stabilizing or affinity-enhancing if they showed negative ΔΔ*G*_bind_ along with folding profiles (ΔΔ*G*_bind_ < 0 kcal/mol and ΔΔ*G*_fold_ < 0.5 kcal/mol). Mutations with a biologically relevant direction of change (stabilizing in MxA and disruptive in NP) were selected based on the magnitude of change or to represent the range of ΔΔ*G*_bind_. Moreover, we also considered the contact frequency of the wild-type residue and the chemical nature of the side-chain substitution for further shortlisting.

### Sequence alignments

Reference sequences of THOV NPs were identified using Blast and downloaded as GenPept files from NCBI. These were aligned with MAFFT, and phylogenies were generated with FastTree in Geneious Prime 2025.1.2. For the MxA alignments, reference sequences of Euarchontoglires were identified with Blast and downloaded as GenPept files, then manually trimmed prior to alignment using MAFFT and FastTree.

### Experimental methods: Plasmid preparation

Site-directed mutagenesis was performed to introduce mutations of interest to pQXCIP-3xflag-MxA and pCAGGS-THOV_NP. Primers for mutagenesis were designed using NEBaseChanger and Snapgene. After PCR, the PCR product was separated on 1% agarose and purified using the Zymoclean Gel DNA Recovery Kit (D4007) from Zymo Research. Plasmid vectors were prepared using FastDigest restriction enzymes and similarly gel-purified. Fragments and vector were then assembled using Gibson Assembly Master Mix (E2611L) from NEB. The Gibson product was then transformed into NEB5a *E.coli* cells, which were plated on LB+Ampicillin agar and incubated overnight at 37°C. Individual colonies were then transferred onto a separate LB+Ampicillin agar plate, and simultaneously inoculated into 5 mL of LB+Amp overnight at 37°C, for 15-18 hours. Plasmids were purified using the PureYield Plasmid Miniprep System. Relevant regions of each NP or MxA gene within each plasmid were sequenced by Sanger sequencing using primer 5’-ATTTCTGGAGAGGCGAAAATGGAAGAAGAAC–3’ for VN04 NP, 5’– CGAGAACCACTTCGTGCTGACGTACCCA– 3’ for THOV NP, or 5’–CGTGGTAGAGAGCTGCC – 3’ for MxA. Plasmids identified as having mutations of interest and consistent flanking sequences were sent for full-plasmid sequencing via Plasmidsaurus. Corresponding colonies were then inoculated into LB+Amp media and grown at 37°overnight, then combined 1:1 with 50% glycerol and stored at -80°C.

### Experimental methods: Minireplicon assay

Minireplicon assays were run in HEK293T cells grown in 96-well black, clear bottom plates from Greiner (catalog no. 655090). HEK293T cells were grown in DMEM with 10% FBS and 1% Penicillin/Streptomycin on treated tissue-culture plates. Cells were grown at 37°C in a humidified incubator with 5% CO_2_. Cells were seeded at a density of between 2.5×10^5^ and 3.5×10^5^ cells/mL, using 100 μL per well, and allowed to grow overnight to between 50-70% confluency prior to transfection. For the THOV minireplicon, 4 ng each of PB2, PB1, PA, and 1 ng of NP, all in pCAGGS vector, along with 20 ng of pHH21-vNP-*FF-luc* (*firefly luciferase*), and 50 ng of pTK-Ren-Luc (*Renilla luciferase*) were transfected into cells along with 50 ng of pQXCIP-3xflag-MxA or an equivalent empty vector plasmid, using TranSIT-293 Transfection Reagent (Sigma MIR 2700). At 24 hours post-transfection, all but 20 μL of cell media were removed from wells, and 20 μL of Dual-glo reagent from the Promega Dual-glo luciferase assay system (E2940) kit was added and incubated for 10 minutes at room temperature prior to reading on the Biotek Cytation3 plate reader. Then, 20 μL of Stop-and-Glo reagent was added per well, and the plate was again incubated at room temperature for 10 minutes prior to reading. The value from the firefly luciferase signal from each well was then normalized by dividing it by the value of the Renilla luciferase signal. For MxA variant analysis, the “fold-restriction” of all samples was calculated by dividing the average normalized empty vector signal by the normalized signal value for each well. For NP variants, the relative activity of the polymerase was calculated by dividing the normalized signal value of each well by the average normalized value of the WT polymerase complex in the absence of MxA. Separately, to assess escape from MxA, the activity of each NP variant in the absence of MxA was then used as the denominator to achieve a normalized value of NP activity in the presence of MxA, which accounts for differences in baseline activity.

### Experimental Methods: Western Blot

Plates from minireplicon assays were sealed with Nunc aluminum seal tape for 96-well plates (Thermo Scientific, 232698) and stored at -80°C. For western blot, plates were thawed on ice and transferred to 200 µL strip tubes. Samples were vortexed and centrifuged for one minute on a quick spin. 30 µL of each sample supernatant was transferred to a new tube. To each of these samples, 6 µL of G Biosciences SDS-PAGE sample loading buffer [6x] (Fisher 786-701) and 1.8 µL of β-Mercaptoethanol were added. Then, AnyKD Mini-protean TGX Stain-free 15-well 15 μL protein gels from Bio-Rad (4568125) were loaded with 10 μL of sample per well, alongside 10 μL of Thermofisher PageRuler Protein Ladder (26619). Gels were run in TGS buffer for 75 minutes at 100V. Protein was then transferred onto membranes using Trans-blot turbo mini 0.2 mm Nitrocellulose transfer pack (Bio-Rad, 1704158). Membranes were blocked in 1:1 Intercept PBS blocking buffer (LICOR 927-70003) in PBS at room temperature for 1 hour on a rocker. After a brief rinse in 1xTBS (Biorad, 1706435) + 0.1%Tween-20 (TBS-T), the blot was then stained with primary antibody overnight with rocking at 4°C: for NP variant blots, this incubation was with rabbit anti-THOV-NP 1:200 (gift from Laura Graf, Univ. Freiburg); MxA variant blots were incubated with M2 anti-FLAG 1:7500 (Sigma F1804-200UG) and rabbit anti-GAPDH 1:7500 (GTX100118). The next day, all blots were washed in TBS-T 5 times for 5 minutes each, then incubated at room temperature in secondary antibody: 1:10000 goat anti-rabbit IgG-HRP (R&D systems HAF008). After two rinses in TBS-T, blots were incubated in SuperSignal West Pico PLUS Chemiluminescent Substrate (Thermofisher Scientific, PI34577) for 2 minutes in the dark, patted dry with Kim wipes, and imaged. Subsequently, NP variant blots were stripped in Restore PLUS Western Blot Stripping Buffer (Thermo Scientific 46430), rinsed 2x in PBS, then blocked in 1:1 Intercept PBS blocking buffer (LICOR 927-70003) in room temperature, rocking for 1 hour. Blots were then rinsed once in TBS-T, then incubated with additional primary antibodies: M2 anti-FLAG 1:7500 (Sigma F1804-200UG); rabbit anti-GAPDH 1:7500 (GTX100118) for 2 hours on a rocker at room temperature. Blots were then washed 5 times for 5 minutes each in TBS-T, and incubated with secondary antibody (1:10000 goat anti-mouse IgG-HRP (R&D Systems HAF007), 1:10000 goat anti-rabbit IgG-HRP) for 1 hour rocking at room temperature. Blots were then rinsed twice in TBS-T, and again incubated in SuperSignal West PLUS Chemiluminescent Substrate for 2 minutes in the dark, patted dry with Kim wipes, and imaged.

## Acknowledgements

We thank A. Clyde, P. Dietzen, L. Graf, P. A. Rowley, J. Young, and F. M. Ytreberg for comments on the manuscript. We thank L. Graf for the THOV NP antibody. We thank J. Young for the logo plot code in R. We thank M. Wu (Fred Hutch Biostatistics Core) for statistical guidance. Research reported in this publication was supported by the National Institute of Allergy and Infectious Diseases grant U54 AI170792 (principal investigator: N. Krogan) (to H.S.M.), Howard Hughes Medical Institute Investigator award (to H.S.M.), and the National Institute of General Medical Sciences of the National Institutes of Health under Award Number P20GM104420 (to J.S.P.). Computational resources were provided in part by Research Computing and Data Services in the Institute for Interdisciplinary Data Science at the University of Idaho. The content is solely the responsibility of the authors and does not necessarily represent the official views of the National Institutes of Health.

## Supporting Information Figures

**Figure S1**. Domain organization of human MxA and location of the L4 loop.

**Figure S2**. Validation of the AF3-predicted THOV NP model.

**Figure S3**. Structural characterization of the predicted MxA 11-mer binding interface on THOV NP.

**Figure S4**. Homology model of the extended MxA L4 loop bound to THOV NP.

**Figure S5**. Stability of the THOV NP–L4 loop complex during molecular dynamics simulation.

**Figure S6**. Alignments of MxA and THOV NP show the conservation level of sites selected for mutagenesis.

**Figure S7**. Structural comparison of IAV and THOV nucleoproteins.

**Figure S8**. Structural mapping of influenza A virus and JOSV NP residues associated with escape from MxA restriction.

## Supporting Information Datasets

**Dataset S1. In silico single-site mutational scan of the MxA L4 loop using FoldX.** Excel spreadsheet containing ΔΔ*G*_bind_ predictions for all possible single amino acid substitutions at the 40 residues of the MxA L4 loop involved in the interaction interface (40 residues × 19 substitutions).

**Dataset S2. In silico single-site mutational scan of THOV NP using FoldX.** Excel spreadsheet containing ΔΔ*G*_bind_ and ΔΔ*G*_fold_ predictions for all possible single amino acid substitutions at the 62 THOV NP residues located within 10 Å of the MxA L4 loop (62 residues × 19 substitutions).

## Supplemental figures

**Figure S1.**
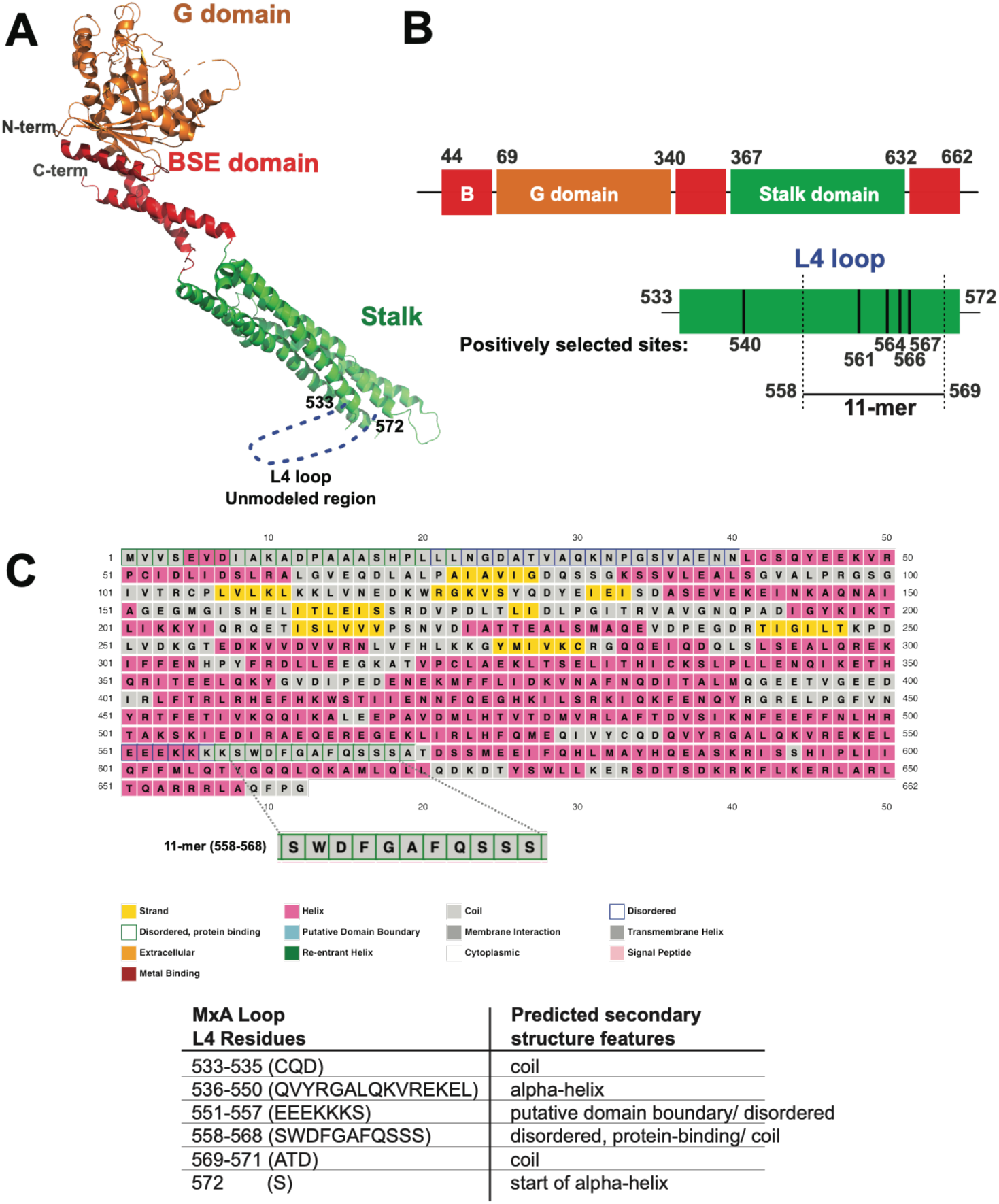
Domain organization of human MxA and location of the L4 loop. **(A)** Crystal structure of human MxA^33^ (PDB: 3SZR) showing the G domain (orange), bundle signaling element (BSE; red), and stalk domain (green). The L4 loop (residues 533–572), which is unresolved in the experimental structure, is indicated by a dashed blue line. **(B)** Schematic representation of the MxA domain architecture. The L4 loop is located within the stalk domain and spans residues 533–572. The 11-mer region first investigated in this study corresponds to residues 558–569. Black bars indicate positively selected sites previously identified in the L4 loop, including residues 561, 564, 566, and 567, as well as residue 540 located upstream of the 11-mer region. **(C)** Predicted secondary structure of MxA and L4 loop (533-572 residues) based on PSIPRED^83^. The predictions highlight the structural and functional features of the region, including the disordered protein-binding segment containing the 11-mer peptide. A summary of the predicted structural elements is provided in the accompanying table.

**Figure S2.**
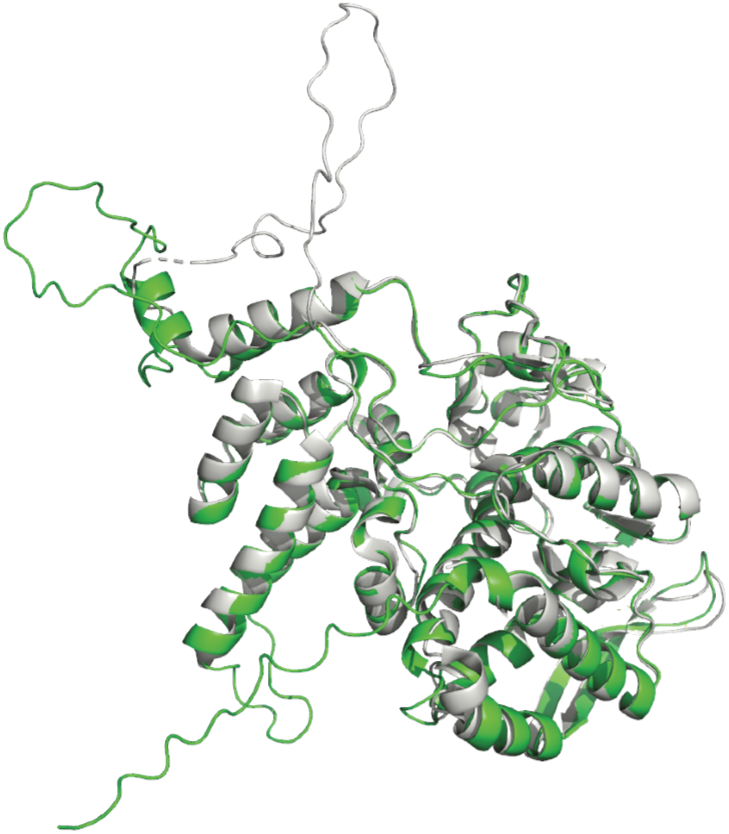
Validation of the AF3-predicted THOV NP model. Structural superposition of the AF3-predicted THOV NP model (green) and the experimentally determined THOV NP crystal structure (gray). The predicted model closely recapitulates the experimental structure, exhibiting high structural agreement (Root Mean Square Deviation < 1 Å).

**Figure S3.**
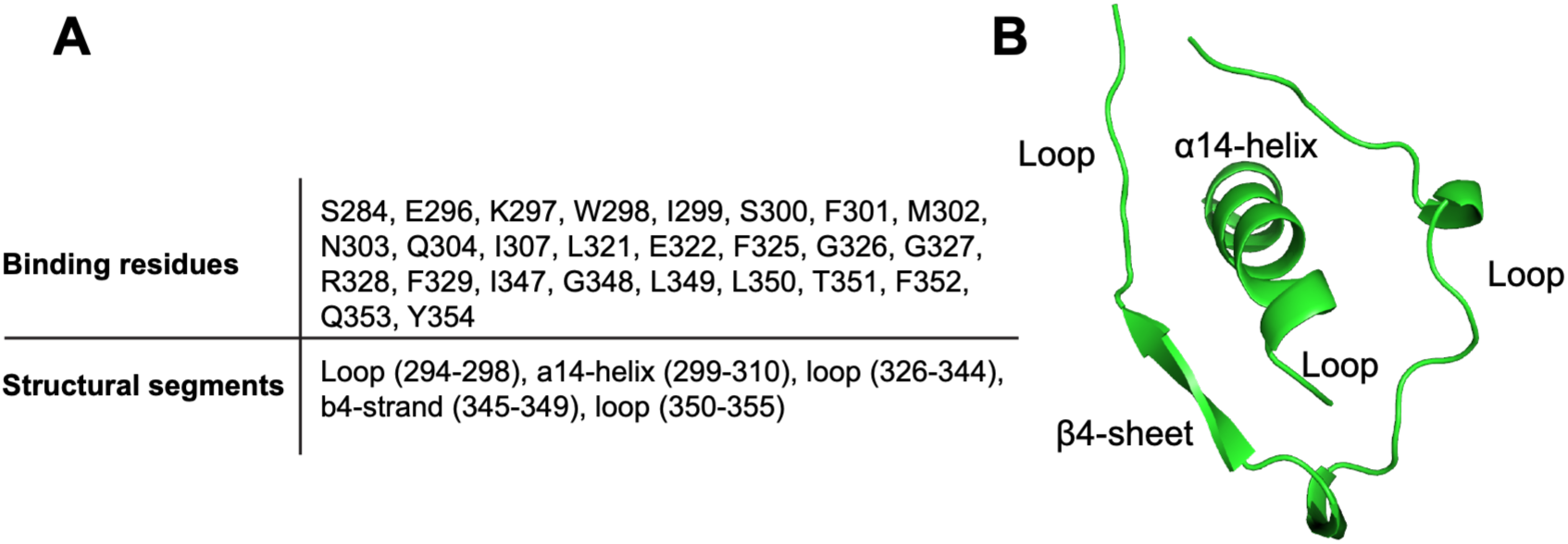
Structural characterization of the predicted MxA 11-mer binding interface on THOV NP. (A) THOV NP residues predicted to interact with the MxA 11-mer peptide and the structural segments that comprise the binding interface. (B) Structural organization of the predicted binding interface. The interacting residues are surface-exposed and located within the NP body domain and can be grouped into five structural segments.

**Figure S4.**
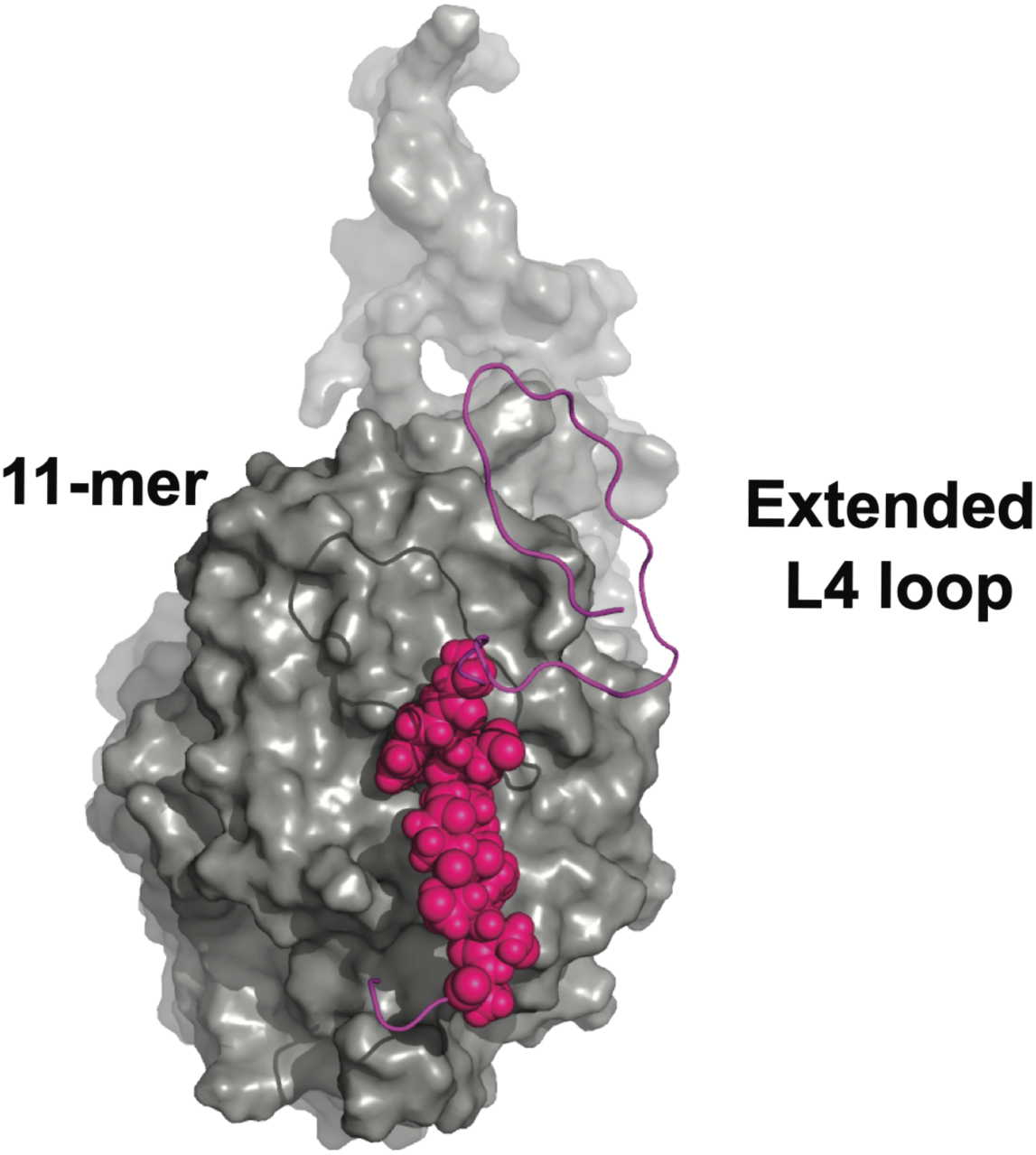
Homology model of the extended MxA L4 loop bound to THOV NP. A homology model of the extended MxA L4 loop (magenta ribbon) bound to THOV NP (gray surface), generated using the top-ranked AF3 11-mer complex as a structural template. The original 11-mer binding region is shown as magenta spheres. Homology modeling was used to reconstruct the full L4 loop while preserving the binding interface and key structural features identified in the previously obtained top-ranked 11-mer AF3 model.

**Figure S5.**
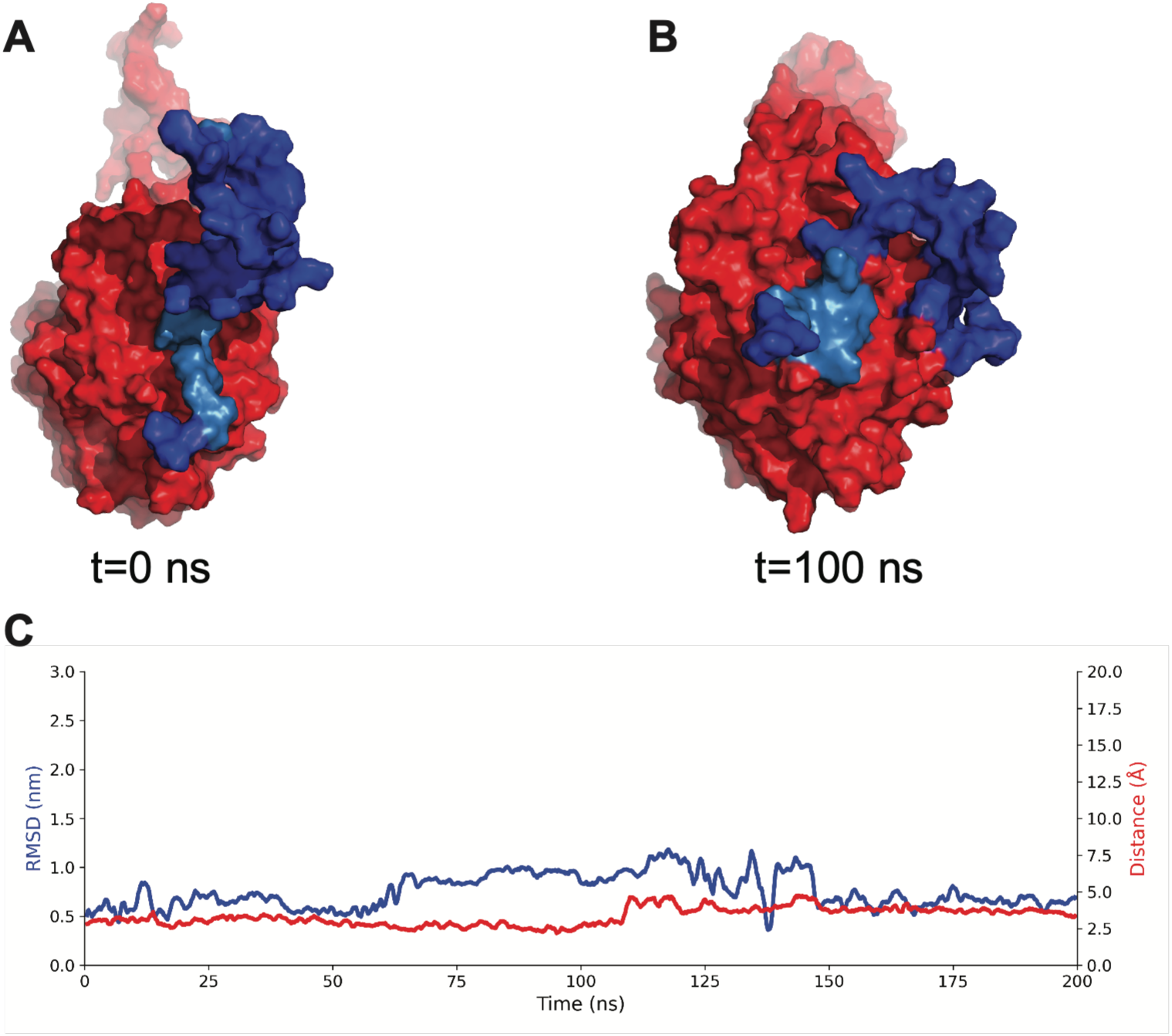
Stability of the THOV NP–L4 loop complex during molecular dynamics simulation. **(A)** Representative structures of the THOV NP–L4 loop complex at the beginning (t = 0 ns) and **(B)** after 100 ns of molecular dynamics simulation. THOV NP is shown as a red surface, the MxA L4 loop as a blue surface, and the 11-mer binding region as a light-blue surface. The L4 loop remains associated with the NP surface while maintaining the orientation of the predicted 11-mer binding interface. **(C)** Time evolution of the complex during a 200 ns simulation. The blue trace shows the RMSD of the MxA L4 loop at each time point relative to the initial structure, while the red curve shows the distance between the 11-mer binding region and the THOV NP binding pocket at each time point. Despite conformational rearrangements of the extended loop, the binding region remains stably associated with the NP surface throughout the simulation.

**Figure S6.**
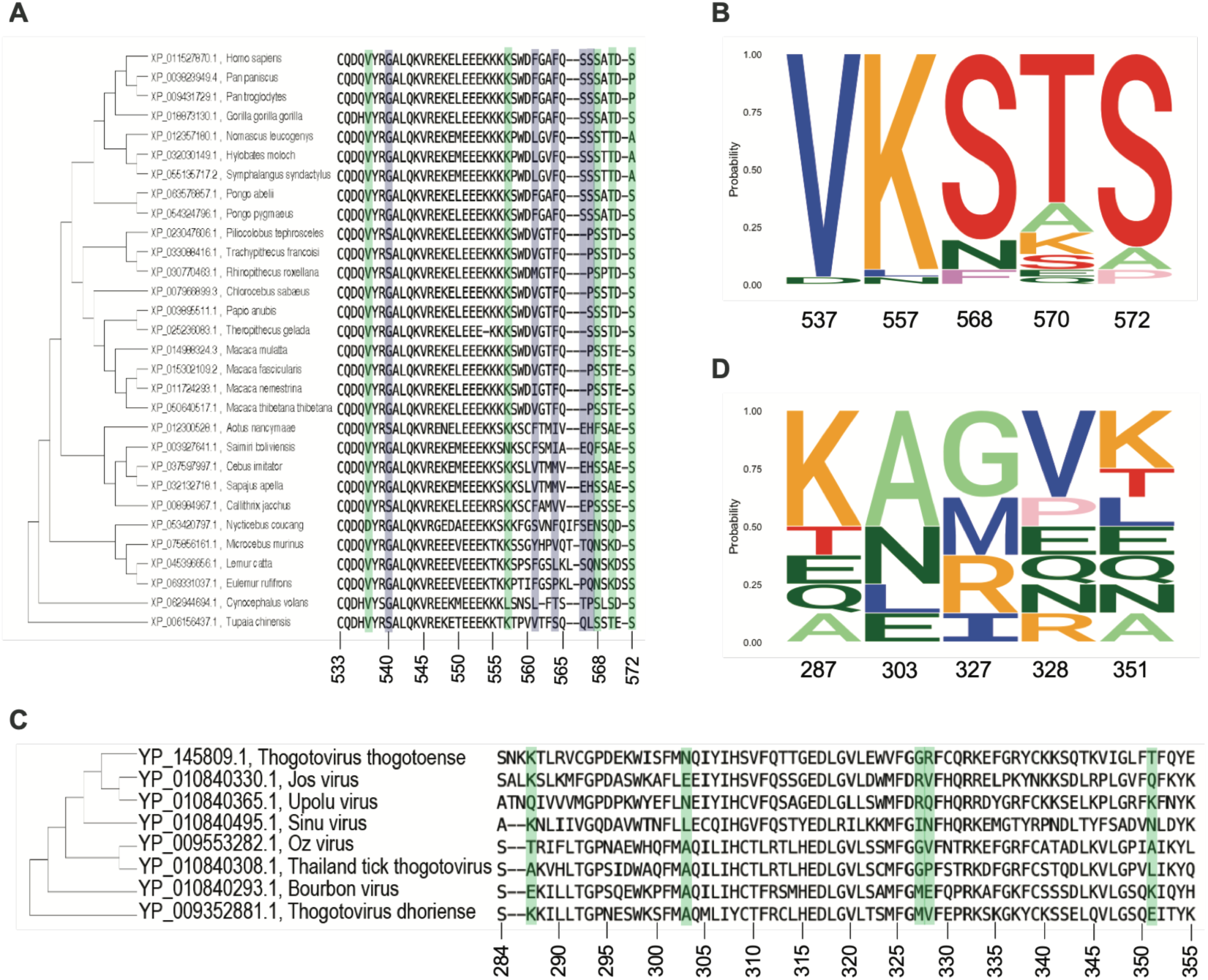
Alignments of MxA and THOV NP show the conservation level of sites selected for mutagenesis. **(A)** Reference sequences of MxA orthologs from *Euarchontoglires* (which includes rodents and primates) were retrieved from NCBI, manually trimmed, and aligned using FastTree and MAFFT in Geneious Prime. The region corresponding to the MxA L4 loop is shown. Positions boxed in green are those selected for mutagenesis in this study. Positions boxed in purple are those previously shown to evolve under positive selection in primates. Positions are labeled at the bottom according to their numbering in human MxA. **(B)** Conservation of residues at each position selected for mutagenesis in the aligned MxA orthologs is represented in a Logo plot. The y-axis represents conservation, with 1.00 representing 100% conservation. **(C)** Reference sequences of Thogotovirus nucleoproteins were retrieved from NCBI and aligned using FastTree and MAFFT in Geneious Prime. Positions selected for mutagenesis are boxed in green. Positions are labeled at the bottom according to their numbering in the *Thogotovirus thogotoense* virus, used in the model. **(D)** Conservation of NP residues selected for mutagenesis among THOV-related viruses is shown in a Logo plot.

**Figure S7.**
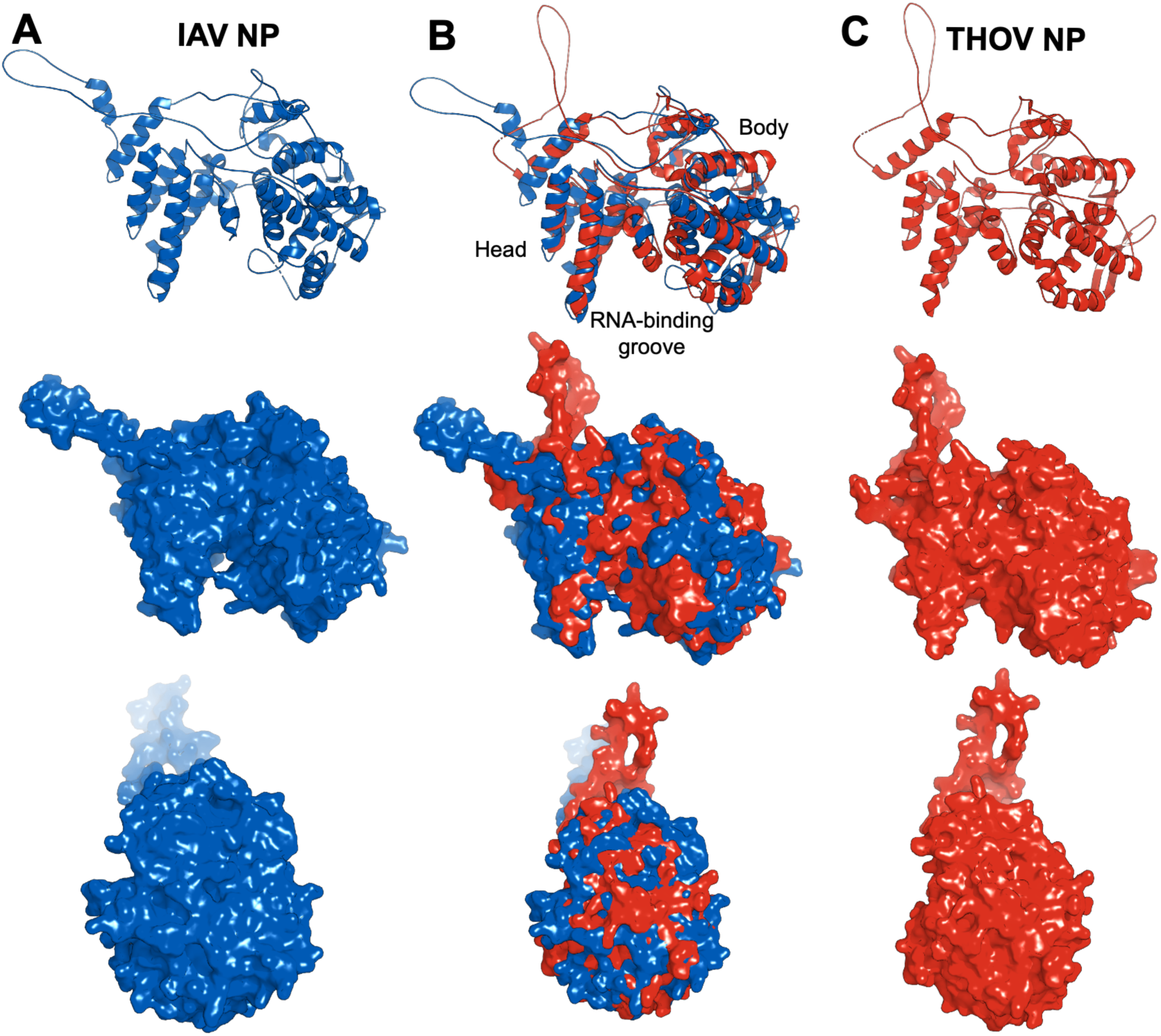
Structural comparison of IAV and THOV nucleoproteins. **(A)** Experimental structure of the influenza A virus nucleoprotein^52^ (IAV NP; blue, PDB ID: 2Q06; 454 residues) shown in cartoon and surface representations. **(B)** Structural superposition of IAV NP (blue) and THOV NP (red). Despite limited sequence similarity, the two proteins share a conserved overall fold comprising a head domain, body domain, and RNA-binding groove. Using Pymol (version 3.1.0)^84,85^, the two NP proteins have an RMSD of 4.03 ÅÅ over 313 Cα-aligned atoms. Using TM-align (version 20220412)^86^, the two NP proteins have an RMSD of 3.68 Å over 409 Cα-aligned atoms. **(C)** Experimental structure of the Thogoto virus nucleoprotein^53^ (THOV NP; red, PDB ID: 8CJW; 467 residues) shown in cartoon and surface representations. The high degree of structural similarity between IAV NP and THOV NP supports the conservation of core nucleoprotein architecture across orthomyxoviruses.

**Figure S8.**
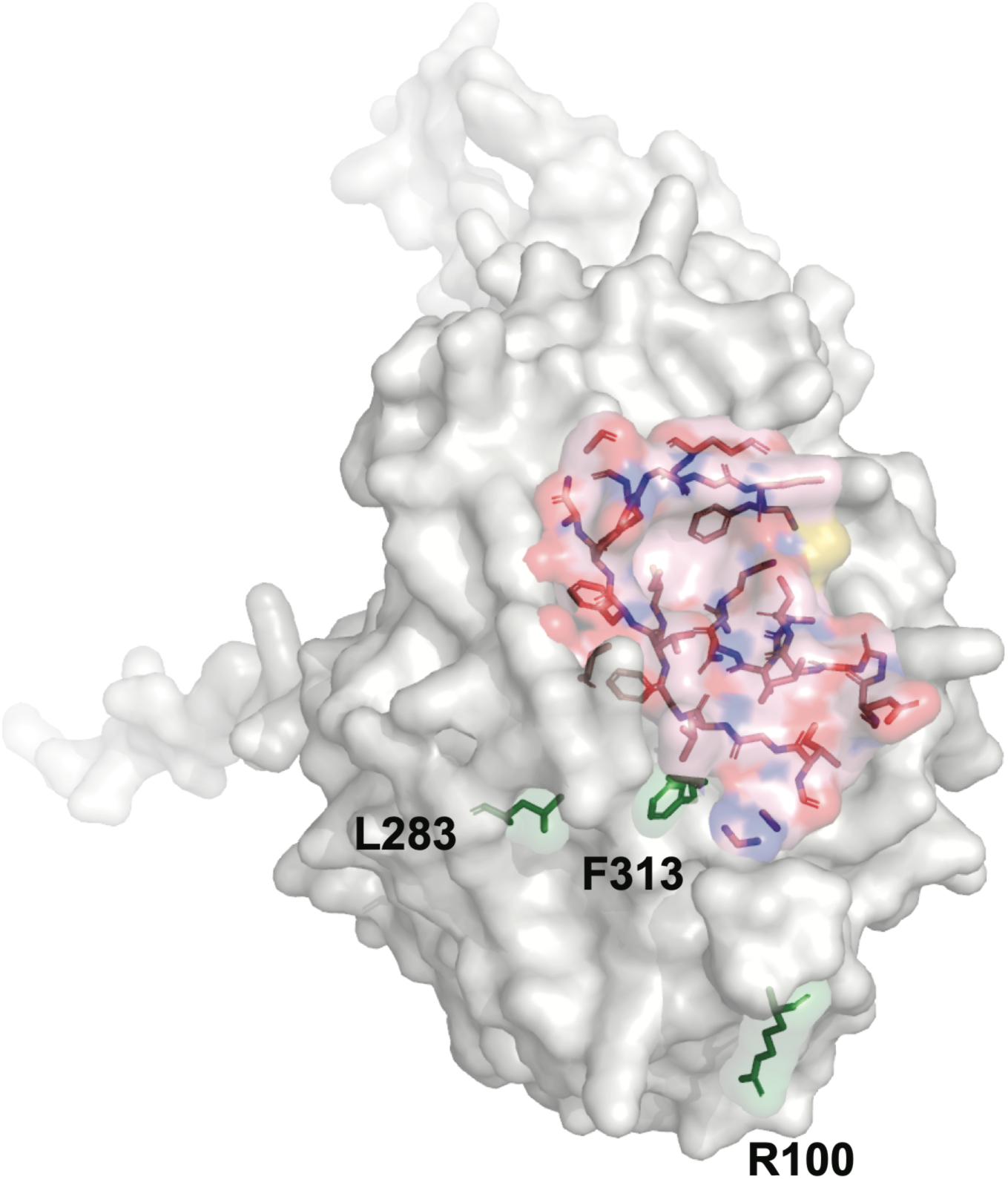
Structural mapping of influenza A virus escape mutations relative to the THOV-MxA binding site. Predicted 11-mer binding site on THOV NP (Figure 2C). Residues predicted to interact with the MxA 11-mer are labeled and colored by physicochemical properties: positively charged (blue), negatively charged (red), and hydrophobic (pink). THOV NP was structurally superimposed with IAV H5N1 NP (IAV NP; PDB ID: 2Q06), and H1N1 IAV residues associated with escape from MxA restriction^46^ are highlighted in green.

